# EASY-CRISPR: a toolbox for high-throughput single-step custom genetic editing in bacteria

**DOI:** 10.1101/2025.03.20.644322

**Authors:** Maxence Lejars, Tomoya Maeda, Maude Guillier

## Abstract

Targeted gene editing can be achieved using CRISPR/Cas9-assisted recombineering. However, high-efficiency editing requires careful optimization for each locus to be modified which can be tedious and time-consuming. In this work, we developed a simple, fast and cheap method for the <u>E</u>diting and <u>A</u>ssembly of <u>SY</u>nthetic operons using CRISPR/Cas9-assisted recombineering (EASY-CRISPR) in *Escherichia coli*. Highly efficient editing of the different constitutive elements of the operons can be achieved by using a set of optimized guide RNAs and single- or double-stranded DNA repair templates carrying relatively short homology arms. This facilitates the construction of multiple genetic tools, including mutant libraries or reporter genes. EASY-CRISPR is also highly modular, as we provide alternative and complementary versions of the operon inserted in three loci which can be edited iteratively and easily combined. As a proof of concept, we report the construction of several fusions with reporter genes confirming known post-transcriptional regulation mechanisms and the construction of saturated and unbiased mutant libraries.

In summary, the EASY-CRISPR system provides a flexible genomic expression platform that can be used both for the understanding of biological processes and as a tool for bioengineering applications.

## Introduction

Gene editing technologies are instrumental for both fundamental and applied research. They provide a way to study the function and the regulation of genes but also to assemble and select novel functions (reviewed in (1)). Classic gene editing methodologies usually rely on the combination of a selectable and counter-selectable-markers with an efficient homologous recombination machinery. In these cases, the induction of an ectopic (often phage derived) homologous recombination machinery is required to achieve high efficiency, in a process called recombineering (2-4). While scarless mutations can be obtained through various approaches using such classic methodologies (5), CRISPR/Cas9-assisted recombineering has been demonstrated to be superior for multiple reasons. First, the *Streptococcus pyogenes* Cas9 endonuclease (Cas9) and a chimeric single guide RNA (sgRNA, fusion of the crRNA and the tracrRNA) are sufficient to induce a targeted double-strand DNA (dsDNA) break upon binding of a perfectly complementary 20 nts sequence (*i*.*e*., spacer of the sgRNA) with the targeted region on the chromosome (6). Hence, an editing sequence, called the repair template, aimed at covering the sgRNA targeted locus, can be used to escape the dsDNA cleavage as well as introduce the wanted mutations. This ensures selectivity of the editing independently of the phenotypic impact of the desired mutation (*i*.*e*., gain or loss of a selectable marker). Second, mutations can be introduced with very high efficiency which enables the construction of saturated libraries of mutants on the chromosome. For example, the CRISPR/Cas9-mediated genomic error-prone editing (CREPE) system can be used for the editing of essential and non-essential genes in *E. coli* with ∼80% editing efficiency (7).

However, the use of such editing approaches is still limited by different factors. First, targeting of a specific locus requires the cloning and testing of candidate sgRNAs to identify suitable ones. Indeed, previous reports suggested that roughly 50% of candidate sgRNAs fail to efficiently kill the recipient strain (8). Furthermore, suitable sgRNAs can also be detrimental and drive an increase in unintended mutations (off-targets) at the genome-scale (9). Second, the system needs to be then introduced in the recipient strain in order to prepare competent cells expressing both the Cas9 protein and the recombineering system before addition of the sgRNA and repair template. Third, editing requires the disruption of the targeted sequence (or the protospacer adjacent motif, PAM). Hence, introduction of scarless mutations is not always possible and often relies on chance to find a suitable target sequence. For these reasons, plasmid systems which are easier to manipulate are often preferred. However, plasmid-borne expression is not perfect as these genetic elements are not very stable and can vary in copy number during growth (10).

To overcome these limitations and make editing easier, we developed the Editing and Assembly of a SYnthetic operon through CRISPR/Cas9-assisted recombineering (EASY-CRISPR) approach. It can be used as an alternative to plasmids to express one (or more) genes, construct reporter fusions and saturated libraries of variants. It allows replacement of open reading frames (ORF) and regulatory elements (promoter and translation initiation region) with low constraints and at high-throughput by targeting a specific locus on the chromosome. The locus to be edited is organized as an operon, facilitating the coordinated expression of multiple genes. Supported by alternative and combinatorial versions of the synthetic operon inserted in three independent loci, the system can be easily upgraded as needed. Moreover, the operon itself can be moved using transduction or recombineering to simplify editing from different loci. Altogether, we now provide an easy and ready-to-use editing tool that would allow most users (experienced or not) to build, test and optimize diverse genetic constructs in *E. coli*.

## Material and methods

### Strain construction, media and reagents

All experiments were performed using an *E. coli* K-12 MG1655 derivative strain, named DJ624, carrying a deletion of the *lacZ* gene (Δ*lac*X74 (11)) and constitutively expressing *lacI* (*mal*::l*acI*^q^). Bacteria were grown in LB medium supplemented with the following reagents (when needed) at the following concentrations, chloramphenicol (25µm/mL), ampicillin (200µg/mL), kanamycin (50µg/mL), zeocine (25µg/mL), isopropyl β-D-1-thiogalactopyranoside (0.25mM), sucrose (10%) and lactose (1%). Strains and plasmids used in this study are listed in Tables S1 and S2 respectively. Their construction is detailed in the supplementary material and methods files.

Oligonucleotides used in this work were ordered from Eurofins Genomics. All PCR amplifications were performed using Q5 polymerase (New England Biolabs) following the manufacturer’s instructions; primer sequences are given in Table S3. Constructed strains and plasmids were systematically verified using Sanger sequencing. Analyzed reporter systems were transduced to a clean DJ624 genetic background using P1 transduction before analysis (as indicated in the strain table).

### Plasmid construction

A sgRNA expression plasmid incorporating the *B. subtilis sacB* gene, encoding a levansucrase, was constructed by combining SS9_RNA (Addgene #716562, (12)) with pk18mobsacB3 (13). The SS9_RNA plasmid backbone (derived from pUC19) was amplified *via* PCR using primers SS9_F and SS9_R. Separately, a DNA fragment containing the *sacB* gene was amplified from pk18mobsacB using primers sacB_F and sacB_R. Both PCR products were assembled using the NEBuilder HiFi DNA Assembly Kit, and the resulting plasmid was named pSS9GR_sacB.

To construct sgRNA expression plasmids, 60 nts ssDNA oligonucleotides, which included the 20 nts spacer region (preceding the PAM site, 5’-NGG-3’) and homology sequences of pSS9GR_sacB were synthesized as 5’-GTCCTAGGTATAATACTAGT-(20 nts spacer)-GTTTTAGAGCTAGAAATAGC-3’. These oligonucleotides, along with a linearized pSS9GR_sacB plasmid prepared via PCR using primers SS9_RNA_F and SS9_RNA_R, were assembled using the NEBuilder HiFi DNA Assembly Kit.

### CRISPR/Cas9-assisted recombineering

Gene editing in the different presented reporter systems was performed as follows. Electrocompetent PAR cells were prepared from exponentially grown cultures (30°C) induced via a 15 min heat shock at 42°C after reaching OD600 0.5. Cells were then quickly cooled in an ice-water bath for 10 min with agitation and washed three times in ice-cold glycerol (10%) before being resuspended in 1/100 volume of ice-cold glycerol (10%). 50 µL aliquots (defined here as a standard reaction, ∼2×10^8^ CFU) can be stored at −80°C for at least 6 months. Editing was performed by thawing an aliquot on ice for at least 15 min followed by addition of 250ng of sgRNA-expressing plasmid and 100ng of repair template. Of note, as repair templates varied in size the absolute number of molecules per reaction varied. In particular, we generally used between 1 to 7×10^14^ molecules per reaction while we only used 6×10^13^ molecules for Ea construction (*lacZ* insertion, Fig. 2A). All components were carefully mixed on ice and incubated for 10 min before electroporation (1.8 kV in 2 mm cuvettes, Fischer Scientific). Cells were immediately resuspended in 1 mL pre-warmed LB medium and agitated in culture tubes for 3h30 before plating on selective medium (ampicillin for selection of the pSS9GR_sacB plasmid and chloramphenicol to ensure the maintenance of the pCREPE plasmid) at 30°C for 24h. Repair templates used for editing were prepared as indicated in the Supplementary material and methods.

### Iterative construction

For iterative constructions, clones obtained after a single round of editing were verified for their phenotype, purified at least once on selective medium and grown in LB with chloramphenicol at 30°C overnight. Cultures were serially diluted and plated on LB in the presence of chloramphenicol and sucrose (10%) before incubation overnight at 30°C. Isolated clones were patched on LB with chloramphenicol versus LB with ampicillin at 30°C to verify curing of the pSS9GR_sacB while retaining the pCREPE plasmid. We typically found that most clones were successfully cured of pSS9GR_sacB. Cells were then prepared for a new round of editing following the protocol indicated in the previous section.

### Gene editing screening and validation

All derivative strains were constructed and validated by Sanger sequencing before testing editing efficiencies. Screening to identify and count recombinant clones was performed based on fluorescence phenotype with the exception of construction Da. Transformants were plated on large Petri dishes (15 cm diameter, Falcon, Corning). CFU were counted using Fiji software (see below) and normalized to total CFU (*i.e*., electroporation of a control reaction without DNA plated only on LB with chloramphenicol for each biological replicate). Total plated reactions were subsampled to verify CFU counting. For constructions Aa, Bb, Cc, Db and Ea, 50 randomly picked clones were patched on selective medium and compared with the purified PAR and validated respective control construction. For the construction Ba, 72 clones were grown in a 96 deep-well plate overnight. Fluorescence was quantified using a plate reader and compared to that of the PAR and validated Ba strain. Some clones (as indicated) were validated using Sanger sequencing. Construction Da (made with repair templates with HA of variable lengths) were validated by PCR (see Supplementary material and methods).

### Construction of libraries of mutant and amplicon preparation

For both *bamA* and *fepA* RBS constructions, three independent libraries each composed of ∼2×10^4^ CFU scraped from a single large plate (50% of a single reaction) were constructed in parallel. Libraries were grown for 5h at 30°C in 200 mL LB medium in presence of selective antibiotics (Amp and Cm) to reduce background noise (*i.e*., to prevent the growth of slow growing escape mutants). Upon reaching OD600 1, cells were harvested. Aliquots were stored at −80°C in glycerol and 2 mL of cells were used for genomic DNA (gDNA) extraction using NucleoSpin Microbial DNA Mini (Macherey-Nagel). Amplicons were PCR amplified from the extracted gDNAs using Q5 polymerase with 20 amplification cycles to prevent over-amplification using primers oML266 and oML267. To control for the accurate representation of the constructed libraries, the oligonucleotides used for construction of the libraries (oML241 and oML243) were used as PCR templates using the same primers and PCR conditions. Of note, primers used to verify the libraries were designed to amplify a region present in both non-edited and edited PAR10 clones (HA) to ensure the detection of non-recombinant clones.

### Sequencing and mutation analysis

#### Sanger sequencing

The sequence of the synthetic operon located in the *argG*-*yhbX* locus was verified by PCR using primers oML203 and oML051 (2133 bp in PAR), the synthetic operon located in the *yfiM*-*kgtP* locus was verified by PCR using primers oML064 and oML065 (2117 bp in PAR5) and the operon located in the *mhpR*-*lacZYA* locus was verified using primers oML266 and oML267 (121 bp in PAR10). PCR products were submitted to Eurofins Genomics and sequencing was performed with at least 2 internal primers (Fw and Rev, Table S3 and see below) in order to overlap the edited fragment. In brief, strain Aa, Ba and Bb were verified using primers oML203 and AK387, strain Ca was verified using primers oML203 and oML049, strains Da and Db were verified with primers oML066 and oML049, strain Ea was verified using primers oML051, oML036, oML186, oML187, oML188 and oML190 to cover the whole *lacZ* gene.

#### Amplicon sequencing for mutant libraries

Amplicons from mutant library constructions (∼120 bp) were gel purified using NucleoSpin Gel and PCR Clean-up (Macherey-Nagel). A second purification step was performed using AMPure XP beads (Beckman Coulter). Quantification was performed using Qubit dsDNA Quantification Assay Kits (Fischer Scientific) and quality control was performed by running samples on agarose gel to control for the integrity of the gDNA. Samples were sent for sequencing (nanopore) using Big Premium PCR sequencing service from the Plasmidsaurus company. Approximately 8000 reads per sample (2 oligonucleotides, 3 *bamA* and 3 *fepA* libraries for a total of 8 samples) were obtained ensuring >200-fold coverage of the library (33 possible sequences including the non-edited PAR10 strain). Conserved regions (between edited and non-edited clones) were used to normalize total reads for sequencing noise (*i.e*., average frequency rate of mutations in the conserved regions). The divergent regions were used to calculate the editing frequency based on the frequency of mismatch. Finally, frequency of G/A at each of the 5 randomized position was calculated for each sample. Of note, this analysis reveals a bias in the distribution of the randomized sequence in the repair templates. This has been confirmed by the manufacturer.

#### Whole genome sequencing

To determine the mutation rate, three independent PAR biological replicates strains (constructed independently by transduction) were edited to construct the Aa, Bb, Ca, Db and Ea derivatives. A total of 15 clones and 3 controls strains were purified twice, grown in 3 mL LB medium at 30°C in selective conditions and their genomic DNA was extracted during the exponential phase of growth (OD600 0.5) using Zymo Quick-DNA Miniprep Plus Kit following the manufacturer instructions (Zymo Research Corporation). A second step of purification was performed using AMPure XP beads (Beckman Coulter). Quantification was performed using Qubit dsDNA Quantification Assay Kits (Fischer Scientific) and quality control was performed by running samples on agarose gel and using a Nanodrop device (Fischer Scientific). Samples were then submitted to the Plasmidsaurus company for hybrid sequencing (long-read using Nanopore and short-read using Illumina technology) and assembly. Variants were called using Breseq (14) by comparing each derivative to its respective parental strain from both short and long reads. In addition, polished assembled genomes (combining both short and long reads) were analyzed using progressive MAUVE alignment (MAUVE software (15) to verify for rearrangements and duplications.

### Fluorescent reporter expression

Following sequencing and validation of the constructed derivatives, edited operons were transduced into a clean DJ624 background by selecting for the antibiotic resistance present on the edited operon before quantification of fluorescence expression (Table S1). Fluorescent protein reporters used in this work display different excitation/emission spectra: sfGFP (488/510), SYFP2 (515/527) and mSca/mScaI3 (569/593). Fluorescence was analyzed on LB plates using a Typhoon FLA9500 (Ex/Em sfGFP: 473/530±10; mScarlet: 532/570±10, Fujifilm) and phase contrast images were captured using an Epson perfection 2450 scanner. Fluorescence kinetics were measured by first growing 3 individually purified clones for each condition in 500 µL LB medium in 96 deep-well plates (Fischer Scientific) for 16 hours at 37 °C with rotary shaking at 1,200 rpm in a Thermomixer C shaker with Thermotop for accurate temperature control (Fischer Scientific). Cultures were diluted 400-fold in 200 µL fresh LB medium, covered with 50 µL sterile mineral oil and grown in flat-bottom black 96-well plates (#655090 Greiner) in a microplate reader (Clario Star Plus, BMG labtech) under continuous orbital shacking (500 rpm) for 16h at 37 °C. Fluorescence (mSca: Ex:560±15, Em:600±15 and sfGFP:Ex:475±10, Em:505±10) and optical density (OD600, 600 nm) were measured every 20 min using enhanced dynamic range to avoid saturation of the signals. Fluorescence was normalized on OD600 and auto-fluorescence was subtracted using non-fluorescent control strains grown on the same plate. Fluorescence values were only compared when obtained from a single microplate.

### Image acquisition and analysis

Fluorescence phenotypes were compared using .tiff images generated from a Typhoon FLA9500 using the Fiji software (16). Contrasts were uniformly adjusted in order to identify strong fluorescence variations in agreement with expected phenotypic variations of the control derivatives and further validated by subsampling and comparison with the control strains on a single plate. CFU counting was achieved in the individual fluorescence channels using the 3D object counter plugin in the Fiji software.

### Sequences for EASY-CRISPR design

Annotated sequences of the different versions of synthetic operons, homology arms, synthetic regulatory elements (promoters, TIR and degron), pCREPE and pSS9GR_sacB plasmids are available as individual open-access files (links provided in Table S4).

Assembled sequenced genomes and amplicons as well as raw sequencing reads (whole-genome sequencing and amplicon sequencing) obtained during this work are available upon reasonable request.

## Results

### Principle of the EASY-CRISPR system

In the EASY-CRISPR system, a synthetic operon, used as an editing platform, can be easily edited thanks to a set of selected sgRNAs targeting different sections of this operon and a CRISPR/Cas9-assisted recombineering module expressed from a plasmid. More precisely, the synthetic operon (Tables S1 and S4) was constructed using recombineering in the intergenic region between the *argG* and *yhbX* genes (Fig. 1A) as used previously ((17) and supplementary material and methods). Two strong bidirectional transcriptional terminators (TT1 and TT2) were added upstream and downstream from the operon to insulate its transcription. The operon is under the control of a constitutively expressed Ptet promoter and it is composed of three cistrons: two fluorescent reporter genes (encoding mScarlet-I, mSca, and superfolder-GFP, sfGFP) under control of two independent and artificial translation initiation regions (TIR1 and TIR2, respectively, Table S4), followed by a kanamycin resistance gene flanked by two flippase recognition target (FRT) sites. In addition, sfGFP is expressed as a fusion with an N-ter flag sequence followed by a linker (*lacZ*57AA: 57 first amino acids of the *lacZ* gene). The kanamycin cassette can be used for transduction of the edited operon and can then be cured as previously reported (3). Alternatively, the upstream *argG* gene can also be used for transduction of the resulting construction into on arginine auxotroph background (as described in (17)).

**Figure 1.**
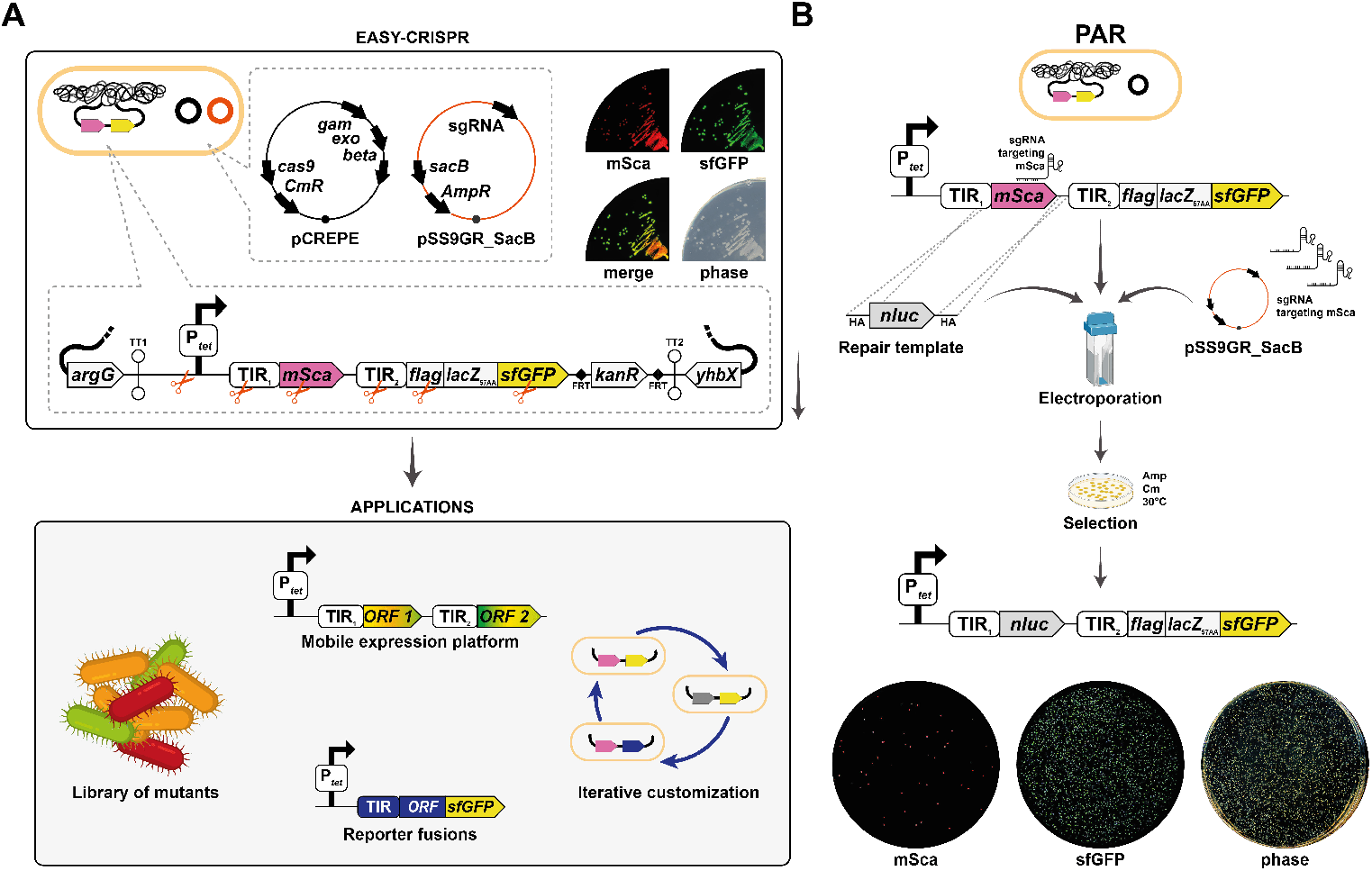
Overview of the EASY-CRISPR system. A) The strain to be edited carries a synthetic operon inserted in the *argG*-*yhbX* intergenic region so that a Ptet promoter drives the expression of two fluorescent proteins (mSca and sfGFP, their individual signals are shown on plates along with the merged signals and phase contrast) and a kanamycin resistance gene (*kanR*). The editing system is divided between two independent plasmids. First, the pCREPE plasmid ensures the constitutive Cas9 expression and the thermo-inducible λ-Red activation. Second the plasmid pSS9GR_sacB allows production of a custom sgRNA targeting one of the regions of the synthetic operon (represented by red scissors). The combined simplicity and high-efficiency of the system make it compatible with various applications, for example as a mobile expression platform, to build libraries of variants or reporter genes. Moreover, operons can be customized using iterative rounds of editing. B) Proof of concept. In this example the parental strain (PAR, sML144) carrying the resident pCREPE plasmid was transformed with the plasmid expressing a sgRNA targeting a region of the mSca gene and a repair template carrying a luminescent reporter (*nluc*). Cells were selected for the presence of the two plasmids. Fluorescence phenotype indicates that the majority of clones were edited, as mSca expression is lost.

The editing system is divided between two plasmids. First, a temperature-sensitive plasmid (named pCREPE, Fig. 1A, black) which ensures the constitutive expression of the Cas9 protein and the thermo-inducible expression of the λ-Red system as previously reported (7). This plasmid can be counter-selected by plating at 37°C. A second plasmid (named pSS9GR_sacB, Fig. 1A, red), introduced by transformation during editing, is responsible for the individual expression of custom sgRNAs, allowing to target different sections of the synthetic operon similarly to what has been previously reported (12). The pSS9GR_sacB plasmid also carries the *sacB* gene, encoding *Bacillus subtilis* levansucrase, enabling the simple and highly-efficient counter-selection of the plasmid by plating on sucrose (13) (see materials and methods).

Classic strategies for gene editing on the chromosome often rely on the selection or counter-selection of a selectable cassette. CRISPR/Cas9 enables the targeting of a specific region of interest to introduce a dsDNA break that is lethal for the bacteria. For this, targeting of a 20 nts sequence (target) located adjacent to a protospacer adjacent motif (PAM, 5’-NGG-3’) on the chromosome is achieved by expression of a chimeric sgRNA which carries 20 nts complementary to the target sequence (crRNA) fused to the tracrRNA ensuring the maturation and loading of the sgRNA (tracrRNA) on the Cas9 nuclease (6).

In the case of the EASY-CRISPR system, we take advantage of sgRNAs specific to the synthetic operon for the complete replacement of the targeted section. Thus, it is not necessary to introduce mutations to disrupt the targeted sequence. In other words, any section of the editable locus can be either deleted or replaced by any DNA sequence of interest. Multiple applications can be foreseen, among which the construction of customized transcriptional and/or translational reporters, libraries of mutants as well as the expression of whole pathways in operons through iterative editing. Moreover, the editing platform is mobile as the operon can be moved to another genetic background using P1 transduction or to another locus using recombineering (see below).

As an illustration of the experimental workflow for editing, the replacement of the fluorescent *mSca* cistron by another gene (*nluc* encoding the nano-luciferase as described in more detail below) is shown in Fig. 1B. An aliquot of electrocompetent cells, kept at −80°C, of the parental strain (PAR, carrying the synthetic operon and the pCREPE, Fig. 1B, top) is transformed with a plasmid expressing a sgRNA targeting *mSca* and a repair template containing the *nluc* gene flanked by homology arms (HA). After plating on selective medium (Amp and Cm at 30°C), thousands of clones are obtained of which the majority have lost mSca expression. It can be emphasized that the method is both inexpensive (only a PCR fragment with short HA is required, see below) and fast as constructions are obtained in one day.

### Identification of suitable sgRNA targeting the editable locus

To target each section of the synthetic operon (Fig. 2A), sgRNA spacers were designed using a prediction tool (https://chopchop.cbu.uib.no/, (18)) with standard parameters. Killing efficiency for each sgRNA was assessed in the PAR strain, in the absence of a repair template, compared with a non-targeting sgRNA (directed against the *nluc* gene, absent from the PAR strain). For simplification, the 5 sections of the operon in the PAR strain were identified by letters (from A to E for Ptet, TIR1, *mSca*, TIR2-*flag*-*lacZ*57AA and *sfGFP* respectively, Fig. 2A). The edited constructions built to assess the efficiency of the system are named with a capital letter, A to E, according to the section that was replaced, followed by a lowercase letter (from a to e) which refers to the construction that was made.

**Figure 2.**
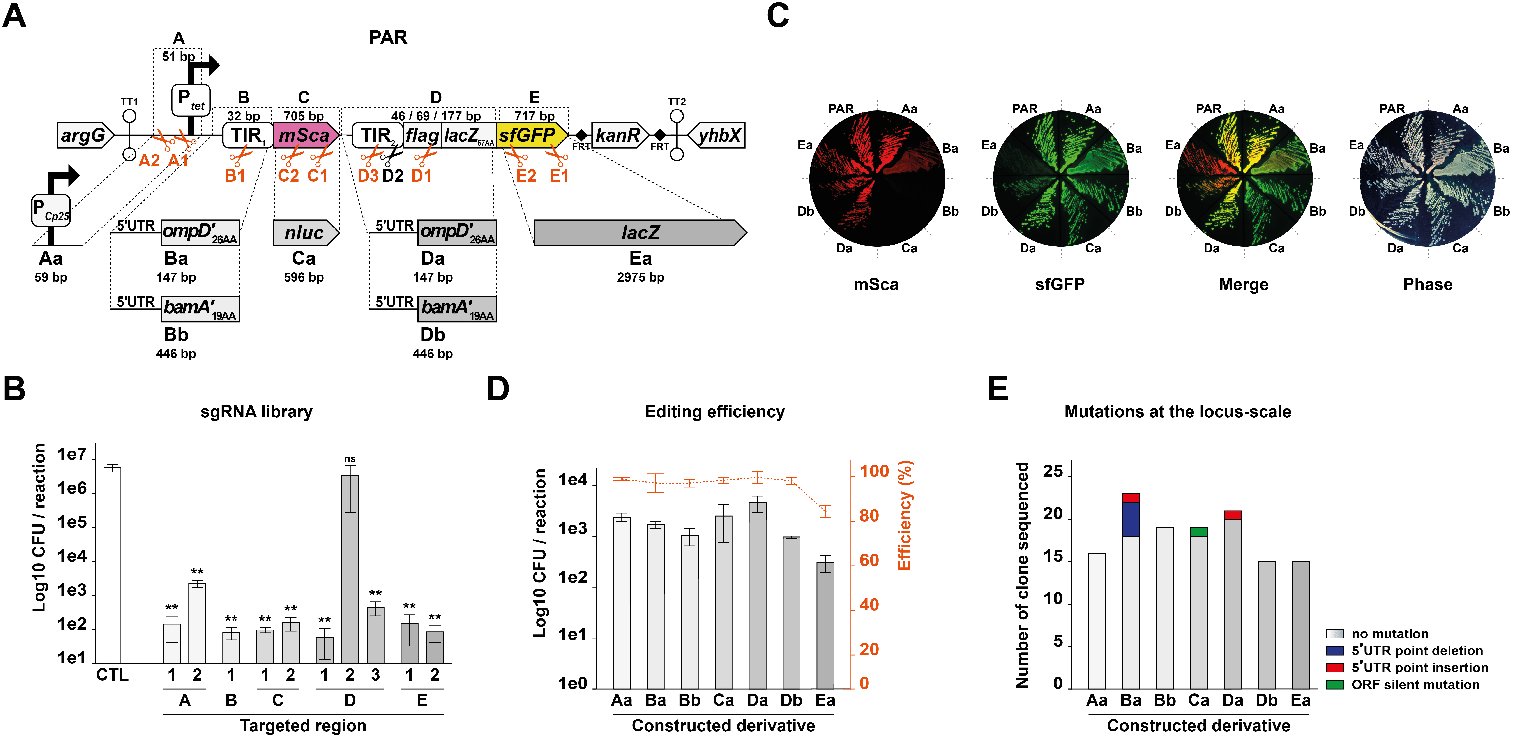
Efficiency and precision of the editing system. A) To test gene edition by EASY-CRISPR, we replaced 5 different sections (A to E) of the synthetic operon carried by the PAR strain, *i.e*., the promoter, the two TIRs and fluorescent reporters, as shown. Scissors represent the positions targeted by the tested sgRNAs numbered according to the region targeted, and shown in red when found suitable for editing or in black when considered not suitable. Throughout the figure, the light to dark grey color-coded boxes (ORF) and histograms bars provides an indication of the region of the operon being considered (light to dark for regions A to E). sgRNA killing efficiency was quantified by counting CFU after electroporation of sgRNA-expressing plasmids in the PAR strain in the absence of a repair template and compared with a non-targeting control sgRNA (CTL). Values (see Table S5A) are means of 3 biological replicates and error bars are standard deviations. Statistical significance was determined by ANOVA (^**^ for p-values ≤0.01). mSca, sfGFP and merged fluorescence signals of the PAR and derivative strains (PAR: sML144, Aa: sML192, Ba: sML153, Bb: sML193, Ca: sML194, Da: sML157, Db: sML195 and Ea: sML196) are shown and compared with phase contrast as a control. B) Total number of CFU and editing efficiency associated with the construction of each derivative. The total number of CFU per reaction (normalized to a standardized reaction containing 2×108 competent cells) is indicated by the histogram bars (y-axis on the left in black). The editing efficiency (dotted lines, red dots, y-axis on the right) was determined based on either fluorescence phenotype or PCR and is indicated as a % of recombinant over total CFU after electroporation of sgRNA-expressing plasmids in the PAR strain in the presence of a repair template. Values (see Table S5B) are means of 3 biological replicates and error bars are standard deviations. C) Mutation rate in the edited locus for each construction was determined using Sanger sequencing on at least 5 clones per biological replicate (*i.e*., 3 biological replicates so at least 15/construction). Clones without mutation are indicated in white/grey (according to the region edited) and fractions carrying point mutations are color coded as indicated (see Table S5C).

We tested a total of 10 sgRNAs targeting the Ptet promoter region (sgRNAs A1 and A2), the TIR1 (B1), the *mSca* ORF (C1 and C2), the TIR2 or N-terminal flag sequence (D1, D2 and D3) or the *sfGFP* ORF (E1 and E2). Their approximate target location is indicated on Fig. 2A. Out of the 10 different sgRNAs tested, 9 were able to reduce the number of colony forming units (CFUs) in a single reaction from 3 up to 5 orders of magnitude (Fig. 2B and Table S5A). Hence, these sgRNA (A1, A2, B1, C1, C2, D1, D3, E1 and E2) were selected for later use in the EASY-CRISPR system. In contrast, the sgRNAs D2 had no effect at all, showing the importance of systematically testing the killing efficiency of a sgRNA.

#### Efficiency of the system to replace the different DNA sections in the PAR strain

We next assessed editing of the synthetic operon carried by the PAR strain. For this, replacement of each of the 5 sections of interest (promoter, TIR1, TIR2-*flag*-*lacZ*57AA, *mSca* or *sfGFP*) was achieved using dsDNA repair templates carrying 51 bp HA. Of note, this length was chosen based on previous reports (2,7). As represented in Fig. 2A, the Ptet promoter (51 bp) was replaced by a stronger PCp25 promoter (59 bp), resulting in strain Aa, constructed using the sgRNA A2. Similarly, the TIR1 (32 bp) was replaced to build *ompD* or *bamA* translational fusions with mSca (147 and 446 bp, strains Ba and Bb, sgRNA B1), *mSca* (705 bp) was replaced with *nluc* (596 bp, strain Ca, sgRNA C1), the TIR2-*flag*-*lacZ*57AA section (292 bp) was replaced to build *ompD* and *bamA* translational fusions with *sfGFP* (147 and 446 bp, strains Da and Db, sgRNA D1) and *sfGFP* (717 bp) was replaced by the *lacZ* gene (2975 bp, from the 58^th^ amino acid, strain Ea, sgRNA E1). Constructions were built using the editing system as described above. Fluorescence of the validated constructions was assessed on plate (Fig. 2C). Where possible screening for replacements was based on large/significant differences in fluorescence levels of mSca or sfGFP and validated by sequencing. It is worth noting that the system does not restrict the editing to replacement of one section by another but more generally permits replacement of any region of the locus by adapting the HA to be used (Table S4).

For each replacement, we typically reached about 10^3^ CFU or above in a single reaction, except for the construction of strain Ea which gave about 10^2^ CFU (Fig. 2D and Fig. S1A). The editing efficiency was estimated based on fluorescence phenotypes from the initial plating (Fig. S1A) and was further validated by patching 50 randomly selected clones that were compared to positive (validated edited strains) and negative (PAR strain) controls (Fig. S1B, Table S5A). In the case of the construction Ba (*ompD-mSca* fusion), the fluorescence is reduced 5-fold compared to the PAR strain as measured from liquid cultures (Fig. S1C). Contrary to what can be observed on purified clones (PAR versus Ba, Fig. 2C), differences in fluorescence were difficult to reliably assess from initial plates where colonies are smaller (Fig. S1A). Hence, 72 clones were grown in liquid medium and their fluorescence was measured from overnight cultures to calculate editing efficiency. This allowed us to quantify the number of edited clones but it also highlighted clones where mSca was not expressed (Fig. S1C) suggesting the presence of point mutations in the construction (see below). In the case of the *ompD-sfGFP* translational fusion (strain Da), we observed a similar sfGFP signal as the PAR strain (Fig. 2C) and we thus quantified the editing efficiency for this strain *via* a PCR test to identify non-recombinant clones in pools of clones (Fig. S1D).

Overall, when considering mean results obtained from three independent biological replicates, we found that for all 7 constructions with the exception of Ea, editing efficiency was between 95 to 98% (Fig. 2D and Table S5B). In the case of construction Ea, where the whole *lacZ* gene replaces the sfGFP reporter, editing efficiency decreased to 83% while the effective number of edited clones was one order of magnitude lower than for the other editing events (see above). This difference could be due to the large size of the insert (2975 bp) a known limitation for CRISPR/Cas9-assisted recombineering technologies (19). In brief, replacement of any of the 5 tested sections, using dsDNA repair templates carrying 51 bp HA (as indicated in Table S4) can be achieved at an efficiency comprised between 83% and 98% and yields a large number of clones per experiment.

#### Precision of the editing

While obtaining a high number of edited clones is important, it is also crucial to ensure that the editing is precise. To verify this, we used bidirectional Sanger sequencing on the synthetic operon locus from at least 5 clones from each biological triplicate of each construction (*i.e*., a total of ≥15 clones/construction). Sanger sequencing revealed the presence of point mutations in a minority of clones (6 out of a total of 126 clones analyzed, Fig. 2E and Table S5C) for a mutation rate of ∼0.05 mutations/clone. Mutations are mainly found as point deletions in the 5’untranslated region (5’UTR) of fusions and we see a clear bias in the construction of the Ba strain (*ompD*-*mSca* translational fusion, 5 mutations in 23 clones, ∼0.2 mutation rate) which may reflect a lower quality of the repair template or a less favorable recombineering event. This confirms what we observed when analyzing the mSca fluorescence of 72 clones for the Ba construction (Fig. S1C) where ∼17% of clones displayed a strongly reduced mSca expression (12/72). It is noteworthy that we never observe more than 1 mutation/clone and that there is no clear correlation between the distance from the HA and the position of the mutation (Table S5C). Thus, editing of the locus of interest seems relatively precise.

In this work we do not modify the expression of proteins related to the mismatch repair system (*e.g*., overproduction of MutLE32K), which was previously shown to increase the global mutation rate (20). However, CRISPR/Cas9 technology was previously associated with an increase in the mutation rate at the genome-scale (9). We thus verified whole genome mutation rate after a single round of editing by sequencing the genome of strains Aa, Bb, Ca, Db and Ea edited within three independently constructed PAR strains (18 genomes in total). We wanted to analyze both point mutations and larger rearrangement at the genome-wide scale. For this we used hybrid sequencing combining both long and short read (Nanopore and Illumina). We identified 5 independent mutations (4 point mutations and one duplication) from the 15 compared genomes for a total mutation rate of ∼0.3 mutations/genome (Table 1 and Table S5D).

**Table 1:** Whole-genome mutation rate after a single round of editing with the EASY-CRISPR.

| Replicate | Construction |  |  |  |  |
| --- | --- | --- | --- | --- | --- |
|  | Aa | Bb | Ca | Db | Ea |
| 1 | 0 | 0 | 0 | 0 | 0 |
| 2 | 0 | 0 | 0 | 0 | 1 |
| 3 | 0 | 1 | 0 | 2 | 1 |

In summary, the EASY-CRISPR system enables the efficient replacement of any of the 5 tested fragments in the synthetic operon carried by the PAR strain. In addition, our data suggests that editing is very precise at the edited locus and is associated with a relatively low genome-wide average mutation rate.

### Both dsDNA and ssDNA with short HA can be used as repair templates in EASY-CRISPR

With the idea to further increase the ease-of-use of the system, we assessed whether it was possible to decrease the HA length of the dsDNA repair templates. For this, we replaced the TIR2-*flag*-*lacZ*57AA section in the PAR strain with two independent repair templates to construct translational fusions of the genes *ompD* and *bamA* with *sfGFP*, using up and down HA decreasing in length from 100 to 22 bp, (Fig. 3A and 3B). The editing efficiency was calculated for each HA length using the PCR assay or the fluorescence phenotype as described above for strains Da and Db, respectively. For both strains, the editing efficiency was between 94 to 100% for HA lengths ranging from 100 to 44 bp while dropping to 0% for 22 bp HA (Fig. 3A and 3B; Fig. S2A, S2B, S2C and S2D; Table S6A and S6B). Hence, we demonstrate that the system can be used with dsDNA repair templates flanked by HA as short as 44 bp without affecting the editing efficiency which remains close to 100%. Furthermore, consistent with the previously established limitations associated with λ-Red alone (2,3), reducing the size of the HA from 100 to 44 bp does not impact the total number of CFU, while shorter HA of 22 bp led to a strong decrease in CFU (Fig. 3A and 3B).

**Figure 3.**
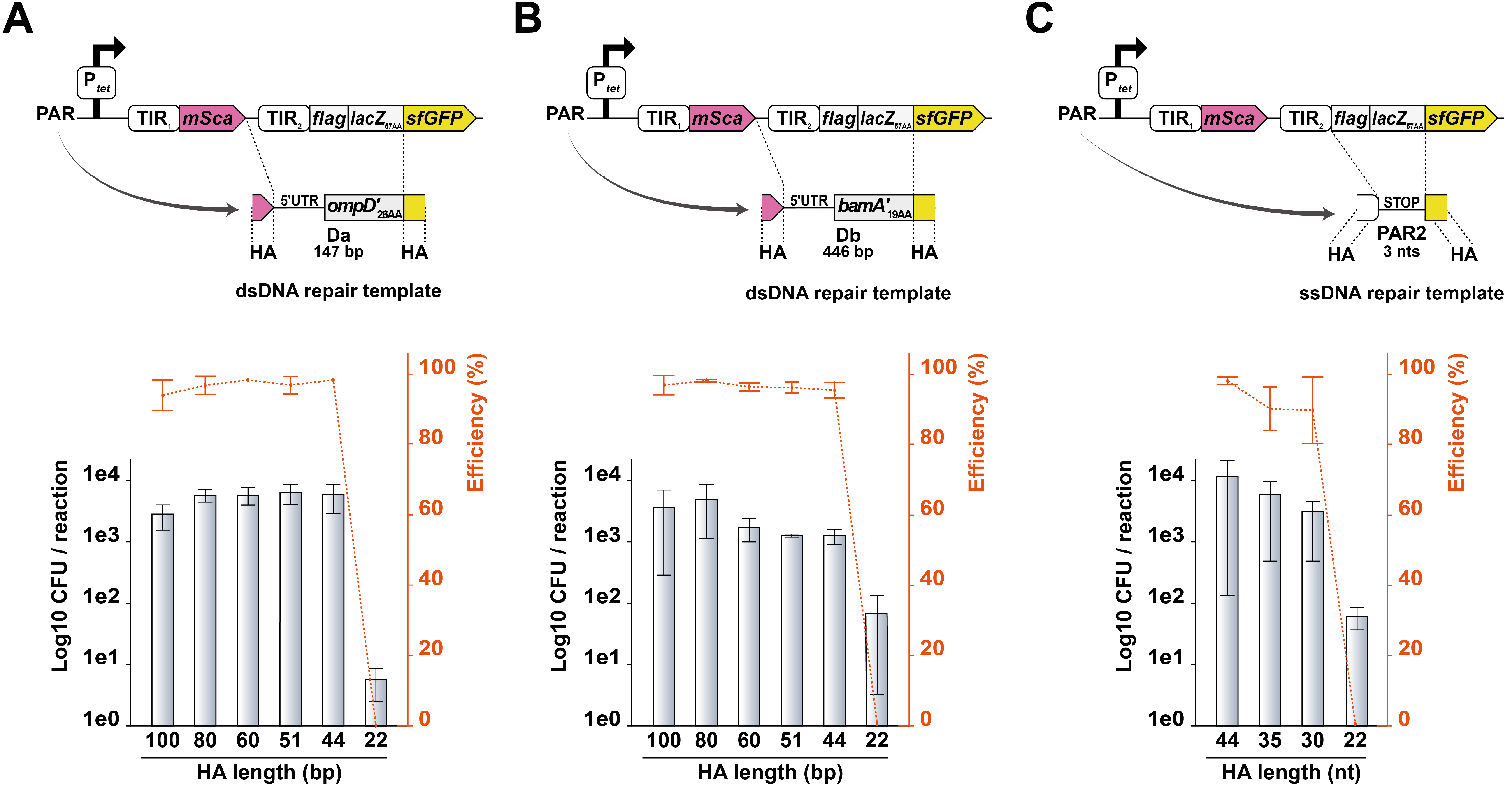
Efficient editing using double and single stranded DNA with short homology arms. The impact of the length of the upstream and downstream homologous arms (HA) on editing efficiency was assessed by electroporation of D1 sgRNA-expressing plasmid and different repair templates: either dsDNA carrying HA ranging from 100 to 22 bp (A and B) or ssDNA carrying HA ranging from 44 to 22 nts (C). For dsDNA repair templates, two independent constructions replacing the TIR2-*flag*-*lacZ*57AA section were tested in parallel with similar HA length variations: (A) strain Da (*ompD*-sfGFP translational fusion, strain sML157) and (B) Db (*bamA*-sfGFP translational fusion, strain sML195). C) Editing was assessed by replacement of the *flag*-*lacZ*57AA region in the PAR strain by a stop codon (PAR2, strain sML245). For all constructions, the number of CFU is indicated by the histogram bars (y-axis on the left in black) and editing efficiency as red dotted lines (y-axis on the right). Values (see Table S6A, S6B and S6C) are means of 3 biological replicates and error bars are standard deviations.

We next wondered if this system is suitable for oligo-recombineering, which provides multiple advantages as compared with dsDNA based recombineering. First, it eliminates the need for a PCR amplification and enables the use of synthetic commercially available oligonucleotides for editing. Second, these commercial oligos can be designed to carry random nucleotides at desired positions which greatly facilitate the construction of libraries of mutants because it does not rely on error-prone PCR. However, from a practical point of view, in the case of oligo-recombineering, the total length, corresponding to the HA and the region to be inserted during editing is limited by the current manufacturing capacity. For example, standard relatively low-priced oligos can be ordered up to 120/150-mer (depending on the manufacturer). Hence, we tried to reduce HA length to maximize the size of the region of interest that could be inserted. For this, we edited the PAR strain to replace the DNA region corresponding to the *flag*-*lacZ*57AA sequence with a stop codon in front of *sfGFP*, using non-modified, high purity salt-free, commercial oligonucleotides (see material and methods) with HA ranging from 44 to 22 nts. As this construct leads to a complete loss of sfGFP fluorescence signal (Strain PAR2, Fig. 3C), we could estimate the editing efficiency based on fluorescence phenotype (Fig. S2E and Table S6C): it varied from 96 to 88% when HA were decreased from 44 to 30 nts but dropped to 0% when HA decreased to 22 nts. Furthermore, decreasing the HA length down to 30 nts in the ssDNA repair template did not affect the total number of CFU (Fig. 3C), while lowering it to 22 nts led to at least one order of magnitude reduction in the number of CFU reaching similar level to that of the sgRNA alone (Table S5A and S6C). Hence, the EASY-CRISPR system enables oligo-recombineering with HA length down to 30 nts while retaining an editing efficiency of 88% which to our knowledge has not been shown before in previously reported CRISPR/Cas9-assisted oligo-recombineering approaches.

### Alternative synthetic constructions increase the modularity of the system

The synthetic operon inserted in the *argG*-*yhbX* locus of the PAR strain represents a flexible and easily adaptable expression platform to express various genes or build reporter systems. While this operon is sufficient for virtually any derivative construction, we also wanted to provide alternative versions, that could still be targeted with the selected sgRNAs, to increase utility of the system.

First, fluorescence phenotype facilitates the verification of the editing and can be used as a screening tool. We were interested to propose alternatives such as the PAR2 strain mentioned in the previous section (Fig. 3C), which allows to start editing from a non-translated sfGFP construction. We also built a derivative expressing only the sfGFP reporter (PAR3) and an alternative operon in which the order of the two fluorescent proteins is inverted (PAR4) as shown in Fig. 4A (top).

**Figure 4.**
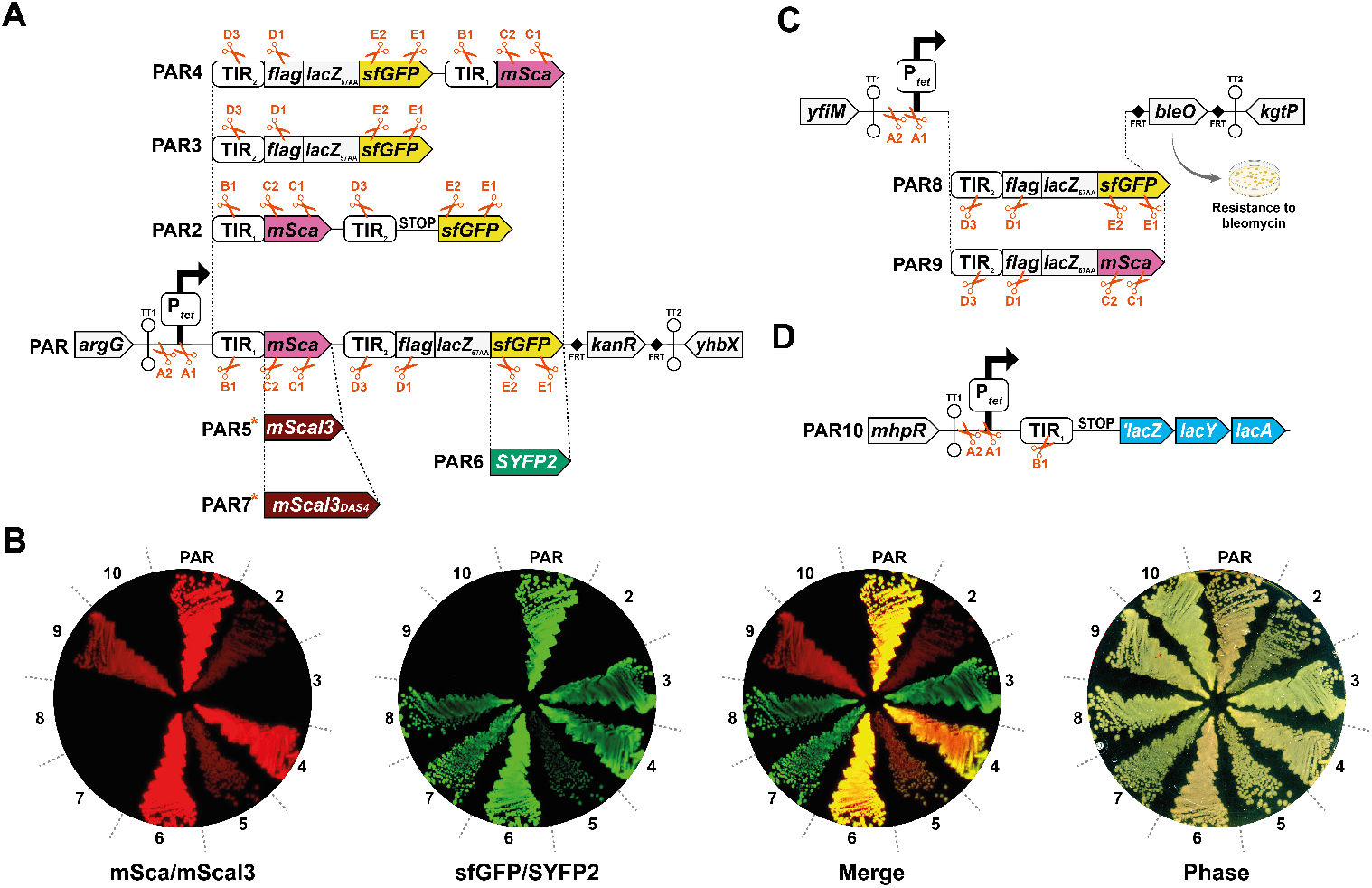
Alternative versions of the PAR strain increase the flexibility of the editing system. A) Schematic representation of the derivative strains expressing respectively mSca followed by a non-translated sfGFP (PAR2, sML245); sfGFP alone (PAR3 sML038); sfGFP followed by mSca (PAR4, sML037); mScaI3 followed by sfGFP (PAR5 sML218); mSca followed by SYFP2 (PAR6, sML166) and mScaI3DAS4 followed by sfGFP (PAR7, sML298). Red scissors represent the positions targeted by sgRNAs suitable for editing. Red asterisks indicate strains that were constructed through two rounds of editing as indicated in the main text. mSca/mScaI3, sfGFP/SYFP2 and merged fluorescence signals are shown for each construction and compared with phase contrast as a control. It should be noted that we observe on plate a significant decrease in fluorescence for mScaI3 as compared to mSca (PAR5 versus PAR) but the difference was not significant when measuring fluorescence from liquid-grown cultures. B) Schematic representation of the alternative synthetic operon, inserted in the *yfiM* and *kgtP* intergenic region, expressing either sfGFP (PAR8, sML299) or mSca (PAR9, sML214) and a bleomycin resistance cassette (*bleO*). Red scissors represent the positions targeted by sgRNAs suitable for editing. C) Schematic representation of the PAR10 strain (sML260) carrying the Ptet and TIR1 fragment followed by a stop codon inserted in between the *mhpR and lacZ* genes. Red scissors represent the positions targeted by sgRNAs suitable for editing.

We also provide alternative versions of the PAR strain expressing fast-maturing fluorescent reporters which could be useful to develop biosensors or biomarkers (21). Both mSca (mScarlet-I, 66 min to reach 90% of the fluorescent signal) and sfGFP (39 min to reach 90% of the fluorescent signal) proteins have been shown to fold and develop fluorescence relatively fast as compared to other reporters (22). However, alternative reporters have recently been found to exhibit similar brightness levels as mSca and sfGFP but with significantly faster fluorescence maturation: in particular, SYFP2 (*a.k.a*. mVenusNB, a fast-maturing variant of mVenus, 18 min to reach 90% of the fluorescent signal) and the recently identified mScarlet-I3 variant (hereafter referred to as mScaI3, estimated to develop fluorescence at least 10 times faster than mSca) (23). These reporters demonstrated a marked improvement in the temporal resolution of time-lapse related experiments. For example, fast-maturing reporters allow the detection of short and transient bursts of expression which is often hidden in fluorescence background noise with classic reporters (22). We thus replaced mSca and sfGFP with mScaI3 and SYFP2, in the PAR strain (to give strains PAR5 and PAR6, Fig. 4A). In addition, we also developed an alternative unstable version of the fast-maturing mScaI3 by addition of an N-terminal degron (24). For this, the DAS4 degron (*i.e*., DAS+4) was fused to mScaI3 resulting in the PAR7 strain (Fig. 4A). As expected, we observe a strong reduction in the fluorescence signal, suggesting that mScaI3DAS4 does not accumulate, as compared with mSca and mScaI3 (PAR7 compared with PAR and PAR5, Fig. 4B). Note that PAR5 and

PAR7 were both constructed through two iterative rounds of editing (PAR edited first to make strain Ca which was then edited to construct strains PAR5 and PAR7, see below). For each alternative construction, in strains PAR to PAR7, at least 6 suitable sgRNAs can be used for editing (Fig. 4A, red scissors).

It is noteworthy that we sometimes observed variations in expression of one fluorescent reporter when the TIR of the second one was modified. This is particularly true in the case of the construction PAR2 were the mSca signal significantly decreases as compared with PAR (Fig. 4B). To a lesser extent, constructions Ba, Bb, Ca, PAR3 and PAR4 also display reduced sfGFP signals as compared to PAR (Fig. 2C and 4B). One possibility is that differences in mRNA structure and/or in the translation efficiency of other cistrons negatively affect the stability of the mRNA (see below).

A major advantage of the system is the possibility to perform iterative editing thanks to the counter-selectable *sacB* marker on the pSS9GR_sacB plasmid and the 9 different sgRNAs able to target the operon. To further illustrate this, strain Ca was cured (see material and methods) from the pSS9GR_sacB after editing while maintaining the pCREPE and competent cells were prepared for a second round of editing. The *nluc* gene in strain Ca was then replaced with mScaI3 or mScaI3DAS4 to build the strain PAR5 and PAR7 respectively, using another sgRNA targeting *nluc* (Fig. 4C) which displays a satisfactory killing efficiency in the strain Ca (2 orders of magnitude less CFU in the absence of a repair template, Table S7). We took advantage of the fluorescent phenotype upon replacement of *nluc* to mScaI3 (construction of PAR5) to verify editing efficiency during a second round of editing. Here again, we obtained ∼10^3^ CFU after editing with an efficiency estimated around 93% (Fig. S3A and Table S7), showing that there is no significant loss in the efficiency with multiple editing steps.

### Efficient editing can be achieved from multiple chromosomal loci

To enable the combination of edited constructions we also provide an alternative version of the synthetic operon which was inserted using recombineering into a different locus of the *E. coli* chromosome: the *yfiM*-*kgtP* intergenic region (Fig. 4C). This locus has been previously shown to be highly transcribed and suitable for ectopic gene expression (25). Derivatives of the synthetic operon at this locus enable the expression of either *sfGFP* (PAR8) or *mSca* (PAR9) and carry complementary sequences for 6 sgRNAs (A1, A2, D1, D3 and E1/E2 or C1/C2 for PAR8 and PAR9 respectively; Fig. 4C, red scissors).

In contrast to the versions located at the *argG-yhbX* locus, we introduced in PAR8 and PAR9 a bleomycin resistance gene (*bleO*). The operon can thus be moved from one background to another as needed, using transduction based on bleomycin resistance selection, or into a *kgtP* deletion mutant followed by selection for growth on α-ketoglutarate that requires a functional *kgtP* gene (26). Editing of these operons could be achieved using sgRNAs (as indicated above) used for editing into the PAR strain. This is supported by previous reports showing that editing of identical sequences from different loci is possible (27). To confirm editing efficiency at this locus, we used oligo-recombineering and the sgRNA D1 to delete simultaneously the fragment containing the promoter, TIR2, *flag*-*lacZ57AA* and the *mSca* gene in the strain PAR9 (Strain DCa, Fig. S3B). As expected, we obtained ∼10^3^ CFU (Table S7) with an editing efficiency estimated at 94%.

We also provide a construction at a third chromosomal site in which the TT1 terminator followed by the Ptet promoter and TIR1 were inserted in the intergenic region between *mhpR* and *lacZ* (PAR10, Fig. 4D). Of note, the 9 first codons of *lacZ* were replaced by a stop codon to prevent translation of *lacZ*. This strain enables the introduction of a fragment of interest in front of the whole *lacZYA* operon which makes it suitable for the construction of individual reporter genes and of libraries of *lacZ* fusions (see the last section of results) combined with screening approaches based on *lac* operon expression (28). Moreover, edited constructs at the *lacZ* locus can be easily moved to a *lacZ* deletion mutant (provided that their expression reaches a sufficient level for Lac+ selection). Hence, PAR8, PAR9 and PAR10 can be used similarly to the PAR strain to build reporters or genetic constructs of interest and then be assembled in the same strain *via* transduction.

In summary, we provide a toolbox of synthetic operon variants that can be easily moved from one locus to another, efficiently edited using the same set of sgRNAs and combined by transduction as needed. This further suggests that, if needed, synthetic operons reported here can be introduced and edited from other loci on the chromosome.

### Application of the tandem-reporter operon to the study of post-transcriptional control

To illustrate a possible application of the system, we designed reporter fusions to study gene regulation at the post-transcriptional level. Post-transcriptional regulation relies on two major mechanisms: those affecting primarily translation or mRNA stability, as represented in Figure 5A (for a focused review see (29)). Typically, transcriptional reporters are designed to provide an estimation of promoter activity or transcript levels by fusion of a promoter or region of interest in front of a reporter gene carrying independent translation signals. In contrast, translational fusions provide information on the level of translation through the fusion of a TIR of a gene of interest to a reporter gene ORF. We thus asked whether the operon organization of the editing system could be used to both quantify the impact and determine the primary mode of action of a post-transcriptional regulator.

**Figure 5.**
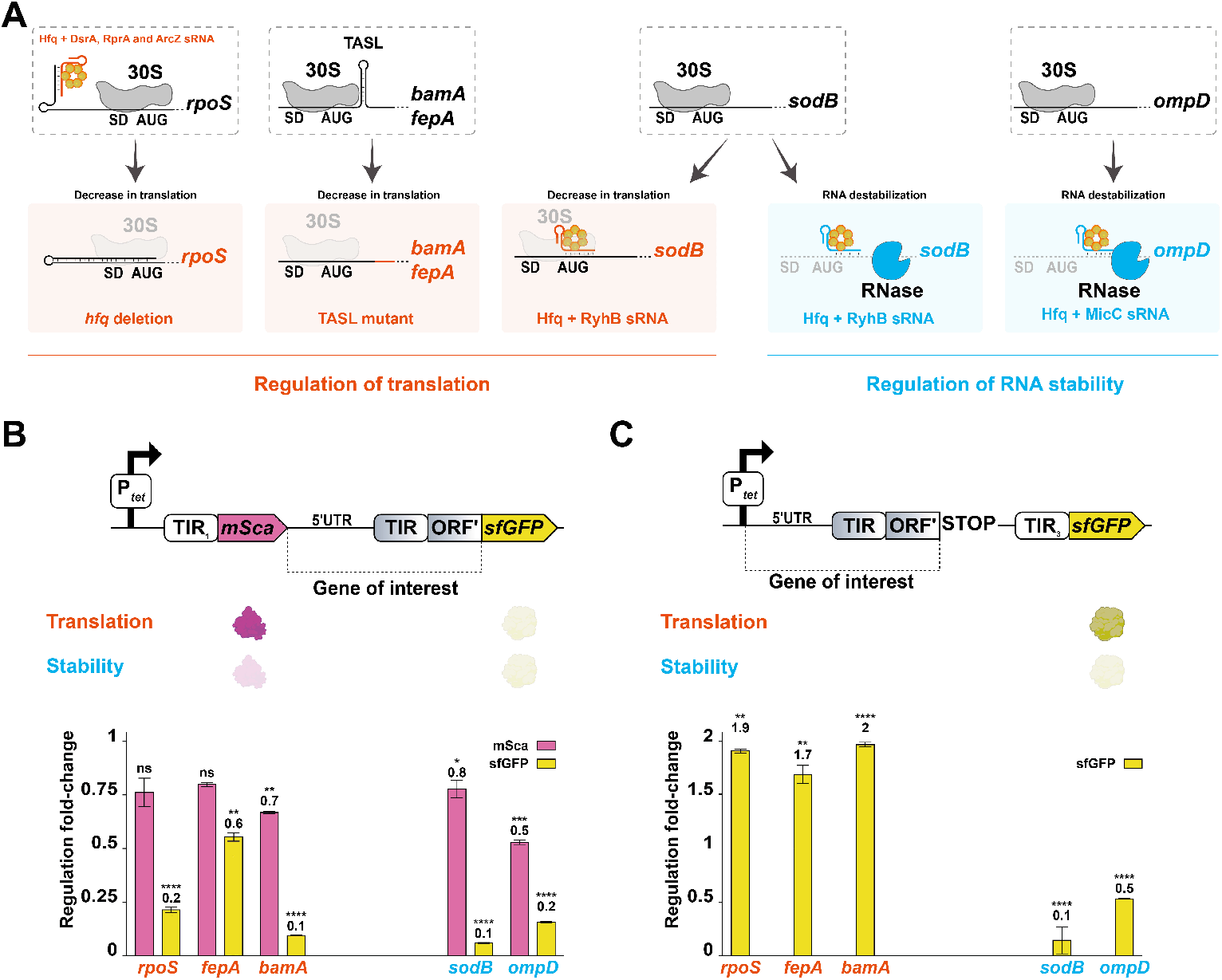
Application of the system to design specific reporter of 5 genes of interest. A) Schematic representation of post-transcriptional regulatory events previously reported to primarily affect translation (left, in red) or mRNA stability (right, in blue). The examples of regulatory events affecting primarily translation tested in this work are the decrease of *rpoS* expression in an *hfq* deleted strain and the inhibition of the mRNA translation of *bamA* and *fepA via* the “mutL” mutations located in the left part of their TASL, as represented in Fig. S4F. Examples of regulatory events affecting mRNA stability tested in this work are the repression of *sodB* by overproduced sRNA RhyB and of *ompD* the overproduction of sRNA MicC respectively. Of note, the RyhB sRNA also represses *sodB* translation (see main text). 30S subunit of the ribosome are represented in light grey when translation is weak and dark grey when translation is strong; mRNAs are represented by black lines, sRNAs are represented in red or blue according to their primary impact on translation or stability respectively, Hfq is represented in orange and ribonucleases (RNase) in blue. B) Tandem fluorescent reporters are used to simultaneously provide a downstream translational (sfGFP) fusion with the gene of interest and an upstream transcriptional (mSca) reporter while (C) mRNA read-out reporters provide a measure of downstream mRNA stability from the sfGFP transcriptional reporter. The expected outcome for both reporters according to the type of regulatory event is schematized under each construction (mSca schematized in pink and sfGFP in yellow with light to dark to indicate low and high expression respectively). Strains carrying tandem fusions (B) used are: *hfq* vs WT for *rpoS*: strains sML281 and sML280; TASL mutant vs WT for *fepA*: strains sML275 and sML274 and *bamA*: strains sML271 and sML270; sRNA expressing plasmid versus control plasmid for *sodB*: strains sML279 and sML278 and *ompD*: strains sML269 and sML268. Strains carrying mRNA read-out fusions (C) used are: *hfq* vs WT for *rpoS*: strains sML295 and sML294; TASL mutant vs WT for *fepA*: strains sML289 and sML288 and *bamA*: strains sML285 and sML284; sRNA expressing plasmid versus control plasmid for *sodB*: strains sML293 and sML292 and *ompD*: strains sML283 and sML282 In all cases regulation fold change is shown relative to the control condition after 16h of growth. IPTG inducer of sRNA expression was added directly in the dilution medium at the beginning of growth. Fluorescence was measured in microplate reader for 16h. Values (Table S8) are means of 3 biological replicates and error bars are standard deviations. Statistical significance was determined by ANOVA (ns for p-values ≥0.05, ^*^ for p-values ≤0.05, ^**^ for p-values ≤0.01, ^***^ for p-values ≤0.001 and ^****^ for p-values ≤0.0001). Kinetic fluorescence measurements from which these values were extracted are shown in Fig. S4 and S5.

Known examples of post-transcriptional control via a primary effect on translation (Fig. 5A) include the synthesis of the RpoS sigma factor, a major regulator in stationary phase and multiple stress responses (30). The *rpoS* mRNA is mainly transcribed as an mRNA carrying a long 5’UTR whose intramolecular folding inhibits translation initiation (31). This repression is alleviated in the presence of the regulatory sRNAs DsrA, RprA and ArcZ that prevent the formation of the inhibitory structure and is dependent on the RNA-binding protein chaperone Hfq (32,33). Consistent with *rpoS* being mainly regulated at the level of translation initiation, a tandem-reporter system showed that within an operon, an upstream translational fusion of *rpoS* displayed a strongly reduced expression in an *hfq* mutant while the downstream transcriptional reporter was unaffected (34). It was also shown that, in some conditions, the sRNAs DsrA, RprA and ArcZ can play a role in preventing the premature Rho-dependent termination of *rpoS* transcription (35). However, this regulation was not observed when *rpoS* expression was analyzed using a tandem reporter system (34). Another example in which a post-transcriptional event primarily affects translation initiation was identified for the genes *fepA* and *bamA* encoding outer membrane proteins involved in iron acquisition and outer membrane protein assembly, respectively. In both mRNAs, stem-loop structures (hereafter referred to as Translation Activating Stem-Loop: TASL) located within the early coding sequence were found to be critical for optimal translation initiation (36).

Conversely, few clear examples (Fig. 5A) have been identified where sRNAs bind mRNA and primarily trigger mRNA decay. One example concerns the *ompD* mRNA, encoding a porin in *Salmonella enterica*, which was shown to be targeted by the sRNA MicC (whose binding site to *ompD* is conserved from *S. enterica* to *E. coli*), and which in turn leads to the fast destabilization of the *ompD* mRNA by the major endoribonuclease RNase E as the primary regulatory mechanism (37,38). In addition, binding of the sRNA RyhB to the *sodB* mRNA, encoding the superoxide dismutase B, blocks *sodB* translation but was also demonstrated to independently provoke mRNA destabilization (39).

Aiming to distinguish post-transcriptional regulatory events impacting primarily translation or RNA stability, we constructed tandem reporter fusions in which the upstream *mSca* cistron is used as a transcriptional reporter for mRNA stability while a translational fusion to the gene of interest is built downstream with the sfGFP reporter. We expected to observe an effect on the sfGFP signal but no impact on mSca signal upon regulation in the case of a primary impact on translation (Fig. 5B, top), and an effect on both mSca and sfGFP signals in the case of a regulation affecting primarily RNA stability. To test this, such tandem reporters were constructed for these five genes, whose post-transcriptional regulation has been already described, using the previously reported fusion boundaries: *rpoS* (34), *fepA* (36), *sodB* (39) and *ompD* (37) with the exception of *bamA* for which a longer 5’UTR was chosen (−389 nts instead of −107 nts (36), see Supplementary materials and methods).

We analyzed expression of the reporters as follows. The *rpoS* reporter was compared in a WT and an *hfq* mutant background in which *rpoS* expression is decreased. In the case of *fepA* and *bamA*, mutations in the TASL were introduced to reduce or restore translation activation as previously reported (36). Finally, *sodB* and *ompD* reporter expression was measured after overproduction of the sRNA RyhB or MicC, respectively (32,37), compared to the vector control.

At the level of the sfGFP translational fusions, we observed in all cases a significant and expected repression (0.1 to 0.2-fold expression compared to the reference condition in general although for *fepA* it was only 0.6-fold, Fig. 5B, yellow bars). We also observed smaller but still significant reductions in the mSca signal for the *bamA, sodB* and *ompD* reporters (0.7, 0.8 and 0.5-fold respectively, Fig. 5B in pink bars, Fig. S4 and Table S8). Although expected for *sodB* and *ompD*, this was not necessarily expected for *bamA* that is primarily regulated at the translational level. It can be noted that, in the cases of *fepA* and *bamA*, two independent mutants of the TASL and a compensatory mutant in which the folding of the TASL is restored were tested confirming these results (Fig. S4BC and Table S8). Hence, no clear correlation can be established between the mode of repression of the translational fusion and its impact on the upstream transcriptional fusion. And, while the tandem reporter works efficiently as a translational reporter, it fails to reliably separate the different types of regulatory events (at least on the basis of previously established mode of action of the tested candidates).

Taking advantage of the EASY-CRISPR system, we thus tested a different design to assess the effects of regulators based on a read-out of mRNA levels. For this, a truncated version of each gene of interest, followed by an early stop codon and an artificial TIR (TIR3, see Supplementary material and methods) is fused upstream of the *sfGFP* reporter. The aim of this reporter design is to show a negative impact on the transcriptional sfGFP reporter when RNA stability is primarily affected, but no impact when translation is primarily affected (Fig. 5C, top).

Such mRNA read-out reporters were built for the 5 genes of interest and tested as described in the case of the tandem reporters. As expected, *sodB* and *ompD* reporters were significantly repressed, while *rpoS, fepA* and *bamA* ones were not (Fig. 5C; Fig. S5; Table S8). Surprisingly, *rpoS, fepA* and *bamA* reporters were even all significantly upregulated in these conditions (see discussion). Of note, in the cases of *fepA* and *bamA*, two independent mutants of the TASL and a compensatory mutant in which the folding of the TASL is restored were tested and confirmed these results (Fig. S5B and S5C). Hence, we could successfully separate regulatory events in two categories which correspond to the expected regulatory events affecting either RNA stability or translation using this transcriptional reporter system.

In summary, thanks to the modularity of the editing system, we could successfully design reporter systems which recapitulated previously established regulatory events at the post-transcriptional level and allowed to discriminate between regulation mechanisms primarily impacting translation vs. mRNA stability.

### EASY-CRISPR allows construction of saturated libraries of mutants

We next aimed to demonstrate that the system is suitable for saturation mutagenesis by building libraries of mutants. For this purpose, we replaced the TIR1 and stop codon of the PAR10 strain with minimal versions of the ribosome binding sites (RBS) of *bamA* and *fepA* (−29 to +3 and −26 to +3, relative to their respective translation start site, Fig. 6A). Within these RBS, 5 nts were randomized with G or A (2^5^ so 32 possible theoretical sequences) using commercially synthesized oligonucleotides carrying 44 nts HA as repair templates. These nts were chosen around the Shine-Dalgarno (SD) sequence of *fepA* (−10-6), and in the (−12-8) region of *bamA* that does not possess a recognizable SD sequence (Fig. 6A). Compared to the previous experiments presented in this work, competent cells were prepared and used without a freezing step. Large numbers of CFU were recovered (an average of 3.8 and 4.1×10^4^ CFU/reaction for *bamA* and *fepA* respectively, Fig. S6A and Table S9A) after plating on selective medium. Libraries were prepared by scraping CFU corresponding to half of the reaction, and are thus composed of ∼2×10^4^ scraped CFU, ensuring >500-fold coverage of the 32 possible sequences.

**Figure 6.**
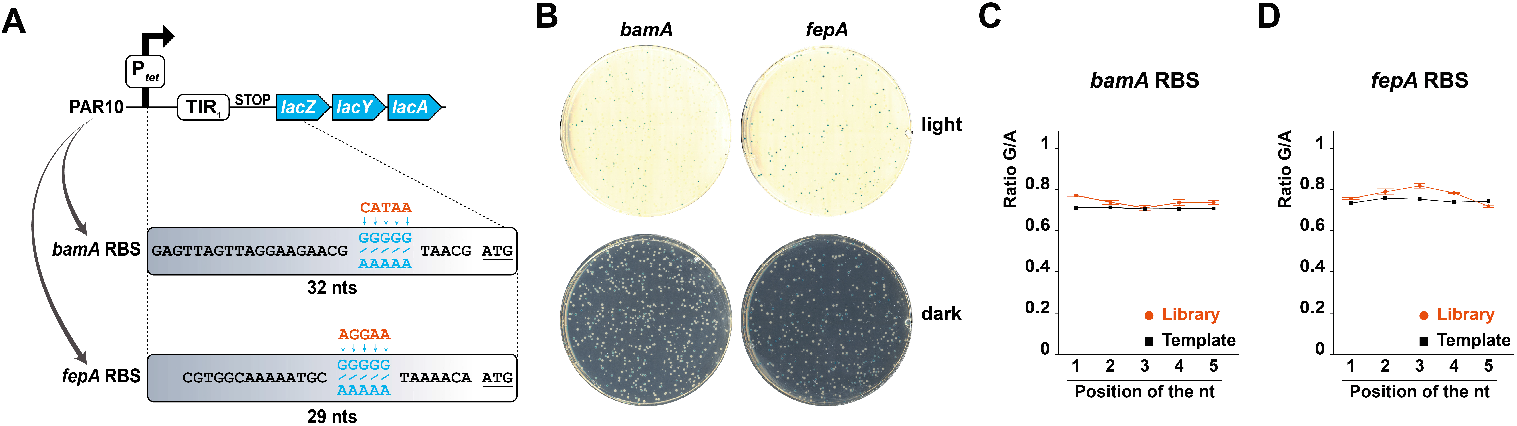
Construction of saturated random libraries of mutants. A) Schematic representation of the PAR10 derivative *bamA-lacZ* and *fepA-lacZ* translational fusions (strains sML296 and sML297) showing a 5 bp randomized (A or G) sequence in the (−10-6) and (−12-8) regions, respectively. The original sequence of *bamA* and *fepA* RBS are indicated in red and in blue the randomized nts that are included in the constructed libraries. Start codons (ATG) are underlined. B) Diluted libraries were plated on LB agar with X-gal and scanned using light and dark background. C and D) Distribution of the randomized nucleotides plotted as a ratio of G/A against the position of the nucleotide (from 5’ to 3’), in black for the repair template and in red for the library for *bamA* (C) and *fepA* (D).

As a first test, we verified the phenotypic diversity among the libraries by plating on medium containing indicators for β-galactosidase activity (X-gal, Fig. 6B and Fig. S6B or MacConkey with lactose, S6B). Libraries show ∼22% and ∼27% of clones expressing *lacZ* at detectable levels for *bamA* and *fepA* respectively (Table S9B). This is a first indication of the diversity of the libraries. To go further, the distribution of mutations in the libraries was analyzed in more detail by deep-sequencing the PCR products amplified from the constructed *bamA* or *fepA* libraries, as well as from the oligonucleotides used as repair templates. Confirming our previous results, editing is highly efficient with 96% and 94% of clones carrying the *bamA* and *fepA* translational fusion respectively (Table S9A). Unexpectedly, we observed a relatively high G/A ratio at each randomized position (∼70% G) in the two libraries (Fig. 6C and 6D). This, in fact corresponds to the distribution of sequences found in each oligonucleotide pool used as the repair template, indicating that the mutant libraries obtained are unbiased and representative of the template material used for editing.

In brief, we demonstrate that the EASY-CRISPR system can be used to construct saturated and representative libraries of mutants in a single step using oligonucleotides with short HA.

## Discussion

In this study, we developed a genetic toolbox allowing the easy and efficient editing of synthetic operons from three loci of the *E. coli* genome using CRISPR/Cas9-assisted recombineering and dedicated sgRNAs. While we mostly have used the construction of reporter genes to provide a validation of the high-efficiency and precision of the system, many other applications can be envisioned. For example, EASY-CRISPR can be used to express one or more genes and coordinate their expression in an operon if needed or can be used to build libraries of variants of genes of interest in a different locus. Moreover, synthetic operons can be easily introduced using transduction in a strain of interest to perform editing in a specific genetic background.

### Advantages and limitations of the system

For a single round of editing, −80°C frozen aliquots of competent cells can be used over a long period of time while ensuring high efficiency. Thus, once the repair templates are available, candidates can be obtained in less than 24h. Repair template design is also facilitated thanks to HA sequences provided in this work (see material and methods). Furthermore, pSS9GR_sacB curing can be easily performed which allows to prepare cells for a second round of editing, when needed, in a short period of time (typically 3 to 4 days). Finally, we provide alternative constructions in 3 different loci (*argG*-*yhbX, yfiM*-*kgtP* and *mhpR*-*lacZ*) which can be combined by transduction. This can be achieved by selecting for the *kanR, bleO, argG, kgtP* or *lacZ* genes as indicated in the results section (Fig. 2A and 4C). Thus, constructions can be combined in a single genetic background. For example, a first step of editing can be performed to build a reporter gene in the *argG*-*yhbX* locus and moved by transduction in a strain carrying a synthetic operon in the *yfiM*-*kgtP* operon to then build a saturated library of variants of a putative regulator and perform screening on the basis of the reporter gene. It should be noted that upon combining synthetic operons in a single genetic context for the purpose of editing, it is crucial to carefully select the targeted section and HA used to ensure that editing can only occur in one synthetic operon.

The EASY-CRISPR system can be used with short HA (44 bp) without reducing editing efficiency. This both facilitates and reduces the cost of production of dsDNA repair templates. Interestingly, 30 bp long HA were previously shown to be sufficient for recombineering using λ-Red alone but at the cost of a reduced efficiency (2). On the other hand, CRISPR/Cas9-assisted oligo-recombineering was thought to require longer HA (60-70bp) to maintain maximal efficiency (40,41). Remarkably, we found that HA as short as 30 nts could be used for high efficiency editing (88%) using ssDNA repair templates. This is advantageous as it allows to reduce synthesis costs but also increases the size of the region of interest that can be inserted using a single oligonucleotide as repair template.

We can also note that some limitations inherent to the system exist. For example, it was previously reported that increasing the size of the insert decreases editing efficiency (19). Indeed, we found a significant drop in the total number of CFU and the editing efficiency when inserting the whole *lacZ* gene as compared to smaller fragments (2975 bp for *lacZ* as compared with 596 bp for *nluc*, Fig. 2D). Partially addressing this limitation, we provide multiple sgRNAs which allow the insertion of large fragments in an iterative manner by replacing one part of a fragment at the time (*e.g*., replace in a first round the N-ter of mSca using sgRNA C2 then in a second round replace the C-ter of mSca using sgRNA C1)).

We show that after a single round of editing, the off-target mutation rate is relatively low (∼0.3 mutations/clone, n=15) as compared with previous reports (*e.g*., ∼1 mutations/clone, n=5 when using λ-Red alone (42)). In this regard, it can be important to move the constructs of interest into the WT or a controlled genetic environment to prevent accumulation of mutations. For this reason, we provide multiple tools to facilitate the transduction of the constructions into another genetic background. Alternatively, it is advisable to work with multiple independent clones after editing.

Moreover, while the targeting of a selected locus by the EASY-CRISPR system ensures its efficiency, we were concerned that it could limit its more general use for other regions. To address this, we verified that the synthetic operon targeted by the EASY-CRISPR can be moved to a different locus and used for editing (*e.g*., at the *yfiM*-*kgtP* or *mhpR-lacZ* locus, Fig. S3B). This strongly suggests that the synthetic operon described in this work or alternative shorter versions can be used to edit endogenous genes or regions of interest. To do this, the operon can be first inserted into the target locus using recombineering. Then, by targeting one of the operon’s sections with a dedicated sgRNA, the operon can be removed while simultaneously introducing mutations into the endogenous gene.

### A tool to study the organization and regulation of gene expression

Illustrating one possible use of the system, we built and tested reporters designed to obtain information about post-transcriptional regulatory events. We first used tandem transcriptional-translational reporters (Fig.5B), and showed that they correctly monitored known effects of the tested regulators or mutations on the sfGFP translational fusions. However, we also found that lowering the translation of the *bamA*-*sfGFP* fusion affected the upstream mSca reporter. One possible explanation is that by reducing *bamA* translation, the TASL mutation could enhance mRNA destabilization as a consequence of the low level of translation. This is consistent with and equivalent to the observed effect of introducing a stop codon upstream of sfGFP (PAR2 construct, Fig. 4A and 4B) that also strongly reduced the mSca signal. Hence, this specific tandem organization does not allow to rigorously discriminate primary effects on translation or stability. It is nonetheless possible that other tandem reporter designs, for instance with a different translational reporter than sfGFP, that can be easily built with the EASY-CRISPR approach, could be optimized for this application.

Indeed, the transcriptional reporter for mRNA levels (Fig. 5C) has successfully recapitulated the mode of action of previously identified regulatory events. Surprisingly, we also found that when used for mRNA readout of the *rpoS, bamA* and *fepA* genes, expression of the transcriptional reporter sfGFP was upregulated approximately two-fold in conditions of expected translational repression (*hfq* deletion mutant for *rpoS*, or *cis*-acting mutations destabilizing the TASL for *fepA* and *bamA* (Fig. 5C). A first possibility is that the close proximity of the upstream stop codon and downstream start codon (26 nts) can result in the occlusion of the downstream RBS. Hence, a decrease in the translation of the upstream cistron can facilitate translation of the second ORF. A second possibility is that the translation of the upstream ORF indirectly impacts the structure of the RNA linker and thus the accessibility of the SD sequence.

### Construction of libraries for high-throughput screening

Thanks to the high efficiency of the EASY-CRISPR system, we could successfully obtain a large number of precisely edited clones. This enables the construction of libraries of variants by using mutagenized repair templates such as error-prone PCR-amplified dsDNA or randomized oligonucleotides. Such libraries are typically used for downstream screening approaches (*e.g*., Deep Mutational Scanning) and for directed evolution approaches (43). We demonstrate that this is possible with the EASY-CRISPR system by constructing two independent libraries (*bamA* and *fepA* RBS sequences, Fig. 6) that are both saturated and unbiased. In particular, we show that using freshly prepared electrocompetent cells we could significantly increase the number of edited clones (∼4×10^4^ CFU/reaction, Fig. S6A and Table S9A) as compared with frozen cells (∼10^4^ CFU/reaction for oligo-recombineering using similar HA length, Fig. 3D).

A saturated library is considered to be sufficient for high-throughput screening when ∼50-fold coverage is achieved (7). Indeed, a screened saturated library provides information not only about gain of function mutations but also about neutral (no bias in distribution between pre-and post-screening) and detrimental mutations (reduced abundance post-screening). Importantly, it also provides quantitative information about the impact of each mutation rather than just qualitative information thanks to large number of mutations for each position. Hence a single mutation library (1 mutation per variant) on 200 bp sequence (800 theoretical possible sequences each containing a single point mutation) would be reached from a single reaction with the EASY-CRISPR system (∼4×10^4^ CFU).

### E. coli and beyond

The EASY-CRISPR system was developed for gene editing in the γ-Proteobacteria *E. coli*, a laboratory workhorse for both fundamental and applied research (11,44,45). Evidence from the literature supports the idea that the system should be easily adaptable to other organisms. First, the λ-Red system was successfully used for recombineering in relatively distant Proteobacteria, for example in the β-Proteobacteria *Burkholderia pseudomallei* or in the α-Proteobacteria *Agrobacterium tumefaciens* (46). Furthermore, CRISPR/Cas9-assisted recombineering was shown to be compatible with multiple bacterial species, relying on λ-Red derived systems or not (47). Hence, previous works support that the EASY-CRISPR could be adapted to other organisms by introduction of the synthetic operon carried by the PAR strain thanks to a selectable marker and by the modification of the two plasmids, pCREPE and pSS9GR_sacB to enable replication in the organism of interest.

## Acknowledgment

We are grateful to F. Quenette for providing *hfq* deletion mutant and to A. Choudhury for kindly providing the pCREPE plasmid and for helpful discussions. We are grateful to members of the group for discussions during this project. We are thankful to J. Plumbridge for critical reading of the manuscript.

