## Supplementary Text and Figures for "EASY-CRISPR: a toolbox for high-throughput single-step custom genetic editing in bacteria"

### Supplementary data

#### Supplementary material and methods

##### Strain construction, media and reagents

Strains carrying synthetic operons described in this study were constructed as derivatives of the strain OK510 (1) carrying a $P_{tet}$-*sacB-cat-'mSca-kanR* operon inserted in the intergenic region between the *argG* and *yhbX* or derivatives of the MG1432 strain (a DJ624 derivative carrying a chromosomal λ-Red system) as detailed in Table S1. In more details, strains PAR3 and PAR4 (Fig. 4A) were first constructed *via* recombineering in strain OK510, as previously described (1). For PAR3, the region of interest was amplified in a first step with primers AK484 and oML019 (Table S3 for list of oligonucleotides) from the pXG30-SF plasmid as a template (2) followed by a second step of PCR using primers AK484 and oML020 to add full-length homology arms (HA). For strain PAR4, the region was amplified using first primers AK484 and oML021 on the pXG30 template and homology arms were added in a second round using primers AK484 and oML022.

The PAR strain was constructed using recombineering in the strain PAR4. For this, a first region ($P_{tet}$-TIR1) was amplified with primers LMO-162a and oML140, using genomic DNA (gDNA) extracted from strain OK510 as a template, followed by a second round of PCR using primers LMO-162b and oML140 to add the TIR1 in front of *mSca*. Full-length HA were then added using two-step PCR. First, using primers oML149 and oML140 and second using primers AK484 and oML140. The final product was recombined into the PAR4 strain. Strains PAR8 and PAR9 were constructed using recombineering in strain MG1432 with dsDNA products assembled using NEBuilder HiFi DNA Assembly Kit (New England Biolabs) from 3 or 5 (for PAR8 and PAR9 respectively) fragments carrying 20 bp overlapping regions. For PAR8 the first fragment (‘*yfiM*-TT1-$P_{tet}$-TIR1-*mSca*) was amplified using primers oML027 and oML032 using gDNA from strain PAR as a template. Fragment 2 (*bleO* gene) was amplified using primer oML033 and oML034 from a dsDNA fragment carrying the bleomycin resistance gene (kind gift of C. Beloin (3)). Fragment 3 (FRT-TT2-*kgtP’*) was amplified using primer oML035 and oML036 from OK510 gDNA. For strain PAR9 the first fragment ((‘*yfiM*-TT1-$P_{tet}$-TIR2-) was amplified using primers oML027 and oML028 from OK510 gDNA. Fragment 2 (TIR2-*flag-lacZ*’) was obtained by annealing primers oML029 and oML030. Fragment 3 (*lacZ*’-*mSca*-FRT) was amplified using primers oML031 and oML032 on OK510 gDNA. Fragments 4 and 5 are the same as fragments 2 and 3 used to build strain PAR8. Strain PAR10 was constructed by recombineering using primer oML254 in the strain MG1508.

It should be noted that the terminator TT1, named ECK120026481, corresponds to the *arcA* terminator and the terminator TT2, named L3S2P21, is synthetic. The efficiency of both TT1 and TT2 was previously established (4). Moreover, TIR1 corresponds to the R148K synthetic ribosome binding site region (5) while the TIR2 corresponds to an artificial ribosome binding site from the pXG30-SF plasmid (2). The TIR3 region (32 bp) was introduced using primer oML206 and corresponds to an intergenic TIR that was successfully used to identify regulatory events affecting primarily stability (6).

All other constructions presented in this study were made using the EASY-CRISPR system and are thus derived from one of the above parental strains. In most cases, derivatives were constructed using a repair template that was amplified in a two-step PCR: a first step to amplify the region of interest (using pairs of forward, Fw, and reverse, Rev, primers) and a second step to complete the full-length HA as indicated in Table S4. For all constructions, with some exceptions as indicated below, gDNA from strain DJ624 was used as a template. For *fepA* fusions, PCR fragments comprising the *lacZ* locus of strains JJ0135, JJ0120, JJ0227 and JJ0180 (7) were used as a template to build WT, mutL, mutR and comp versions of the *fepA* fusions, respectively. For *bamA* fusions, plasmids (unpublished) carrying *bamA* fusions derivative from *lacZ* fusions (strains JJ0383, JJ0384, JJ0385, JJ0386 for WT, mutL, mutR and comp versions respectively (7)) were used. Plasmid derivatives differ in the 5'end of *bamA* fusions which extend to position -389 instead of -107 relative to the translation start of *bamA*. For *ompD* fusions, primers oML071 and oML072 were annealed and used as a template. For *nluc* (8), plasmid pNL1.1 (Promega) was used as a template using primers oML170 and oML171. To replace the $P_{tet}$ promoter with $P_{Cp25}$, amplifying primers (oML179 and oML180) containing the promoter were annealed. For mScarlet-I3 (mScaI3) and SYFP2, plasmid pDx_mScarlet-I3-SYFP2 (Addgene #189763, (9)) was used as a template using primer oML133/oML169 and oML131/174 respectively. For mScarlet-$I3_{DAS4}$ (mScaI$3_{DAS4}$), the optimized degron tag (10) was added using the reverse primer oML233. For *bamA* and *fepA* randomized-SD (RRRRR, with R= G or A), oligonucleotides oML241 and oML243 were used respectively for editing in the PAR10 strain.

#### Gene editing screening and validation

PCR verification of editing efficiencies for strains Da was performed as follows. Isolated clones on plates were picked and mixed in 10µL of water before being boiled for 10 min at 95°C. Pools of cells extracts (5 clones) were obtained by combination of 2µL of each individual boiled sample (total of 10µL) in a new microtube. Pools were tested for the presence of a fragment that can only be amplified in the PAR strain (POS control) and not in the correctly edited strain (NEG control, which shows 2 non-specific weak bands) using primers AK418 and lacZ96 on 1µL of pooled extracts as a template (Fig. S1D). POS and NEG controls were amplified from a pool composed of 1 positive or 1 negative control clone with 4 negative or 4 positive clones respectively to ensure that detection is reliable in a pool of clones. Upon detection of a non-recombinant clone in a pool (presence of a band), each clone from the identified pool was tested individually (Fig. S2B, see gels on the right).

### Supplementary Figures

### Figure S1

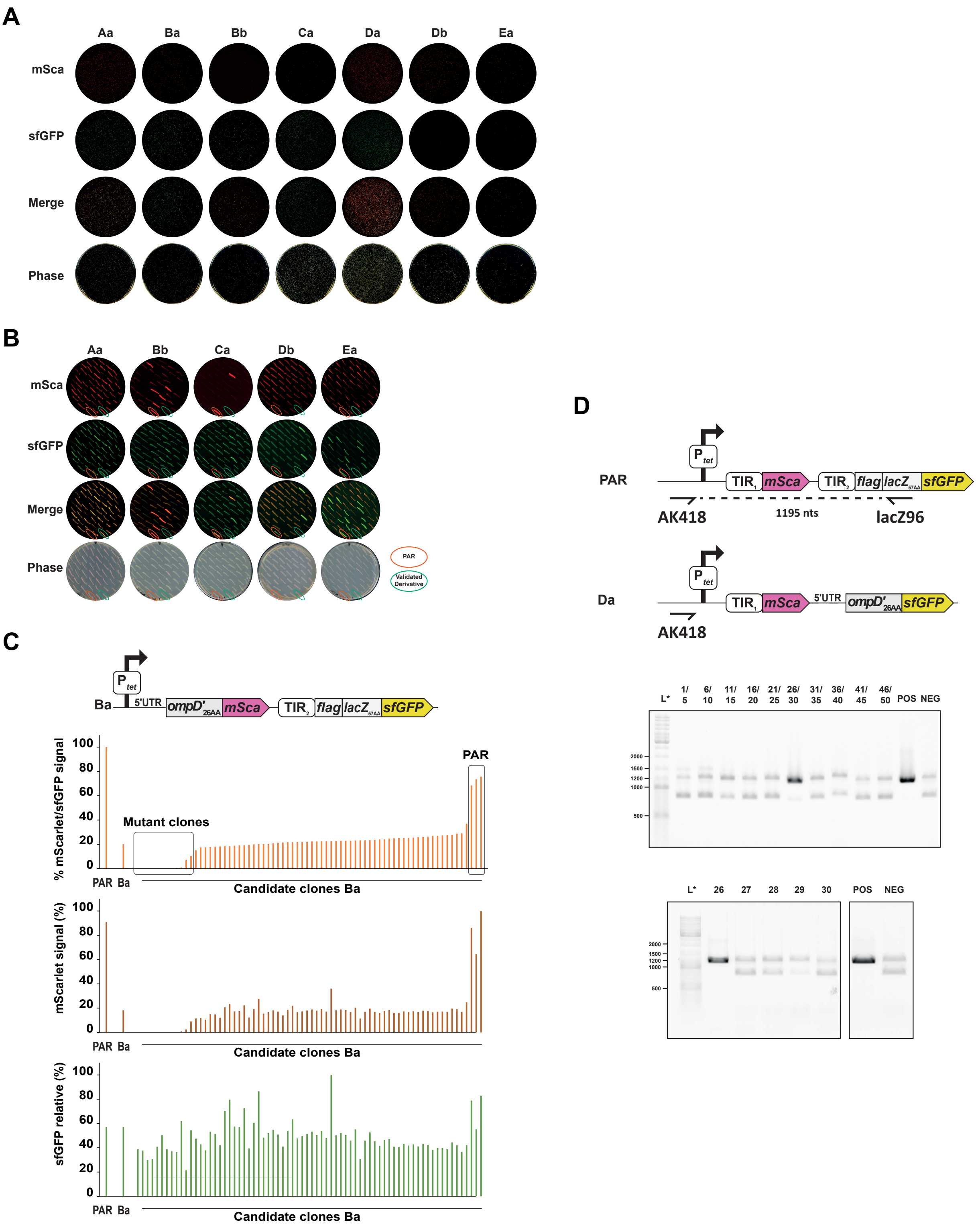


Figure S1. Validation of editing efficiency

A) mSca, sfGFP and merged fluorescence signals are shown for the initial plating on large (15 cm diameter) Petri dishes of editing reactions corresponding to each construction, as indicated in Fig. 2D, and compared with phase contrast as a control.

B) and C) Verification of editing based on phenotype. B) To determine the editing efficiency in the construction of the strains Aa, Bb, Ca, Db and Ea (Aa: sML192, Ba: sML153, Bb: sML193, Ca: sML194, Da: sML157, Db: sML195 and Ea: sML196), 50 clones were tested by patching on selective medium and compared with the PAR strain (red circles) and the corresponding validated strain (green circles). C) For construction of strain Ba, fluorescence of 72 clones was measured from overnight cultures and compared to that of the PAR and validated Ba strains. mSca fluorescence signal was normalized over sfGFP signal and clones were sorted by expression levels. Three clones display normalized fluorescence level corresponding to that of the PAR strain (boxed in black). The 5 rightmost clones were also Sanger sequenced to validate that only the 3 last ones correspond to the non-edited PAR strain. The 12 clones with the lowest mSca fluorescence (square on the left) were also sequenced and found to carry mutations in the ompD-mSca fusion.

D) Verification of editing by PCR. To determine editing efficiency in the Da construction (strain sML157), 50 clones were tested using PCR on pooled clones. Upon detection of a non-recombinant clone in a pool (presence of a band), the 5 clones from the identified pool were tested individually (see gels on the bottom). Regions of complementarity of the PCR primers on the PAR and Da strains are schematically represented, the lacZ96 primer has no complementarity with the Da construction. Two faint bands in all PCR are due to nonspecific amplification.

### Figure S2

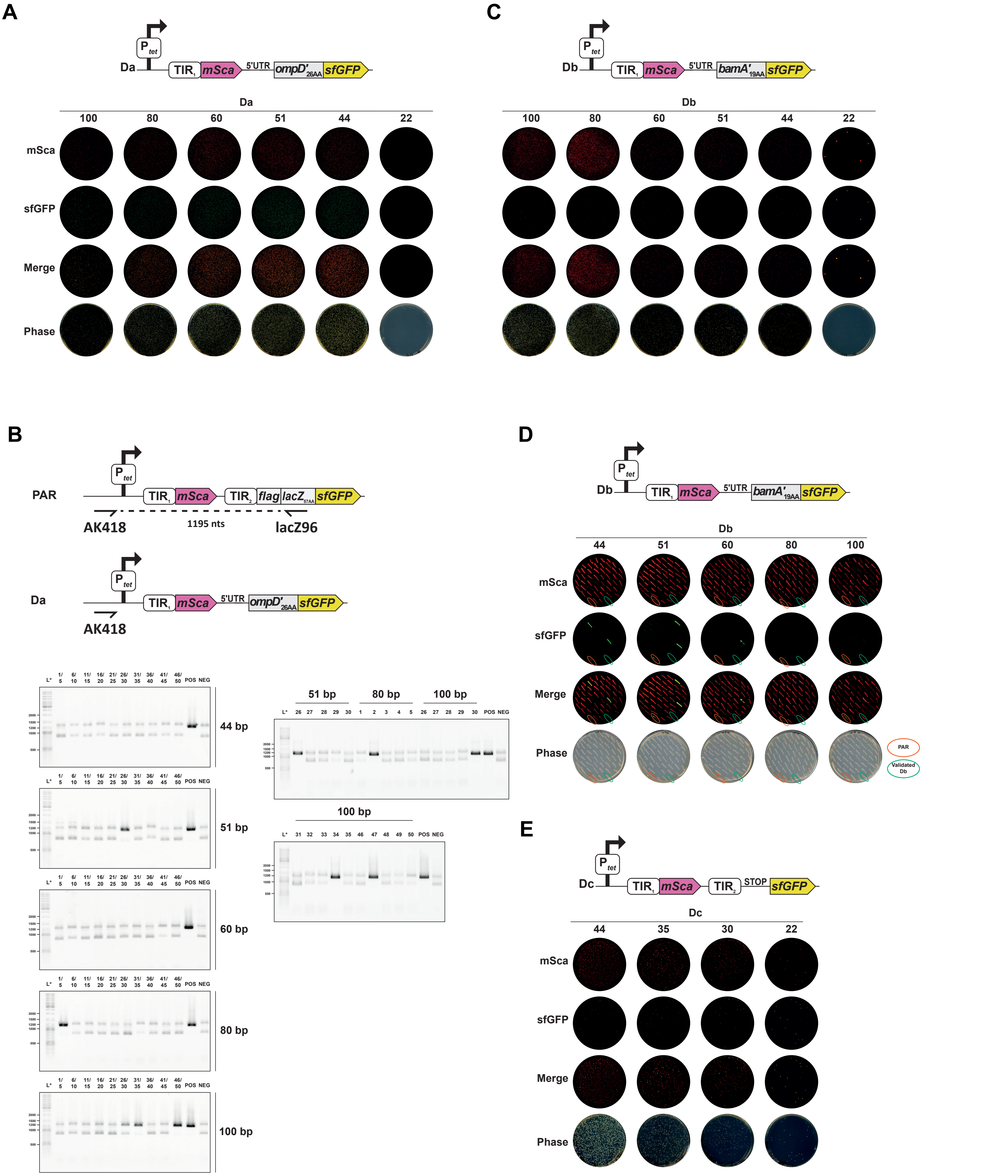


Figure S2. Validation of editing efficiency comparing single and double stranded repair templates

A, C and E) mSca, sfGFP and merged fluorescence signals are shown for the initial plating (on 15 cm diameter plates) of editing reactions corresponding to each construction, as indicated in Fig. 2, and compared with phase contrast as a control. Contrary to panels A and C, only 25% of the reaction was plated in panel E.

B) Verification of editing by PCR. To determine editing efficiency in the Da construction, 50 clones for each homology arm length were tested using PCR with oligonucleotides AK418 and lacZ96 on pooled clones. Upon detection of a non-recombinant clone in a pool (presence of a band in PAR), each clone from the identified pool was tested individually (see gels on the right). Regions of complementarity of the PCR primers (AK418 and lacZ96) on the PAR and Da strains are schematically represented, the lacZ96 primer has no complementarity with the Da construction. Two faint bands in all PCR are due to nonspecific amplification.

D) Verification of editing based on phenotype comparison. To determine the editing efficiency in the Db construction, 50 clones for each homology arm length were tested by patching on selective medium and compared with the PAR strain (red circles) and the correctly edited Db strain (green circles).

### Figure S3

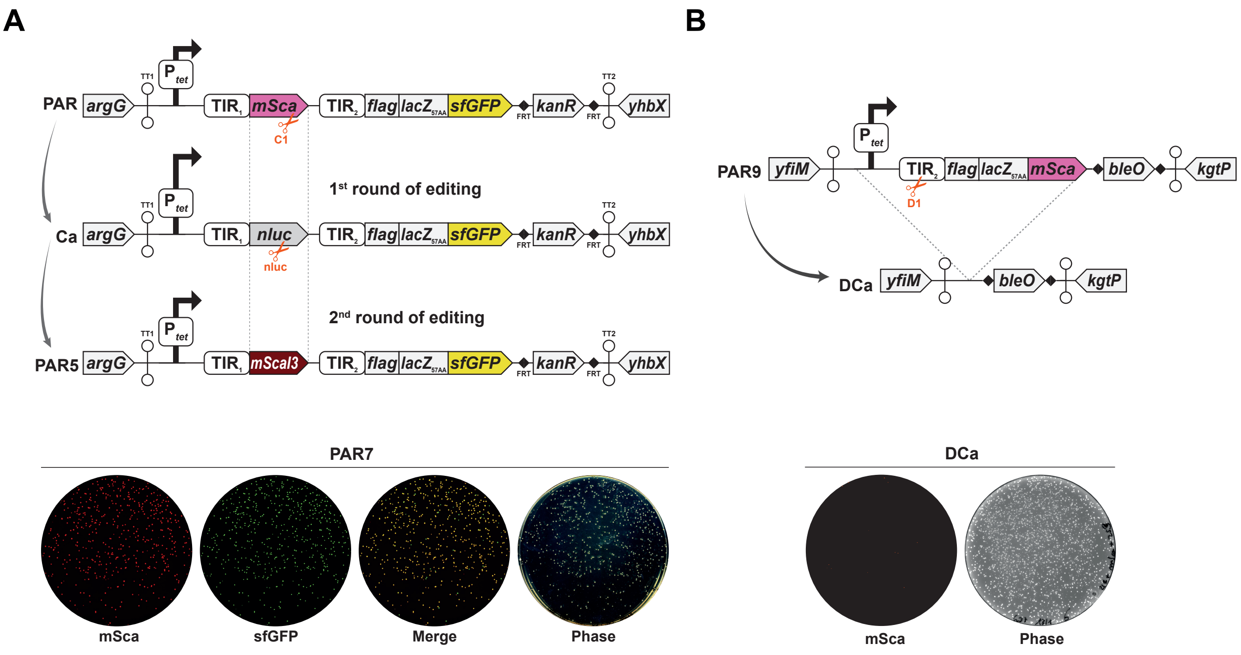


Figure S3. Editing efficiencies in alternative synthetic operons

A) C) Schematic representation of the iterative construction of the strain PAR5 (sML218). To test editing efficiency during a second round of editing, the Ca strain (sML194) was edited using a sgRNA targeting nluc to replace nluc with mScaI3. Total reactions were plated on 15 cm diameter plates, and mSca, sfGFP and merged signals were compared to the phase contrast as a control. Note that strain PAR7 (mScaI3DAS4, sML298) was constructed using the same strategy (see Supplementary material and methods).

B) To test editing efficiency from the yfiM-kgtP locus, the PAR9 strain was edited using the sgRNA D1 to remove the fragment located in between the upstream terminator and the bleO gene. Total reactions were plated, the mSca signal compared to the phase contrast control is shown.

### Figure S4

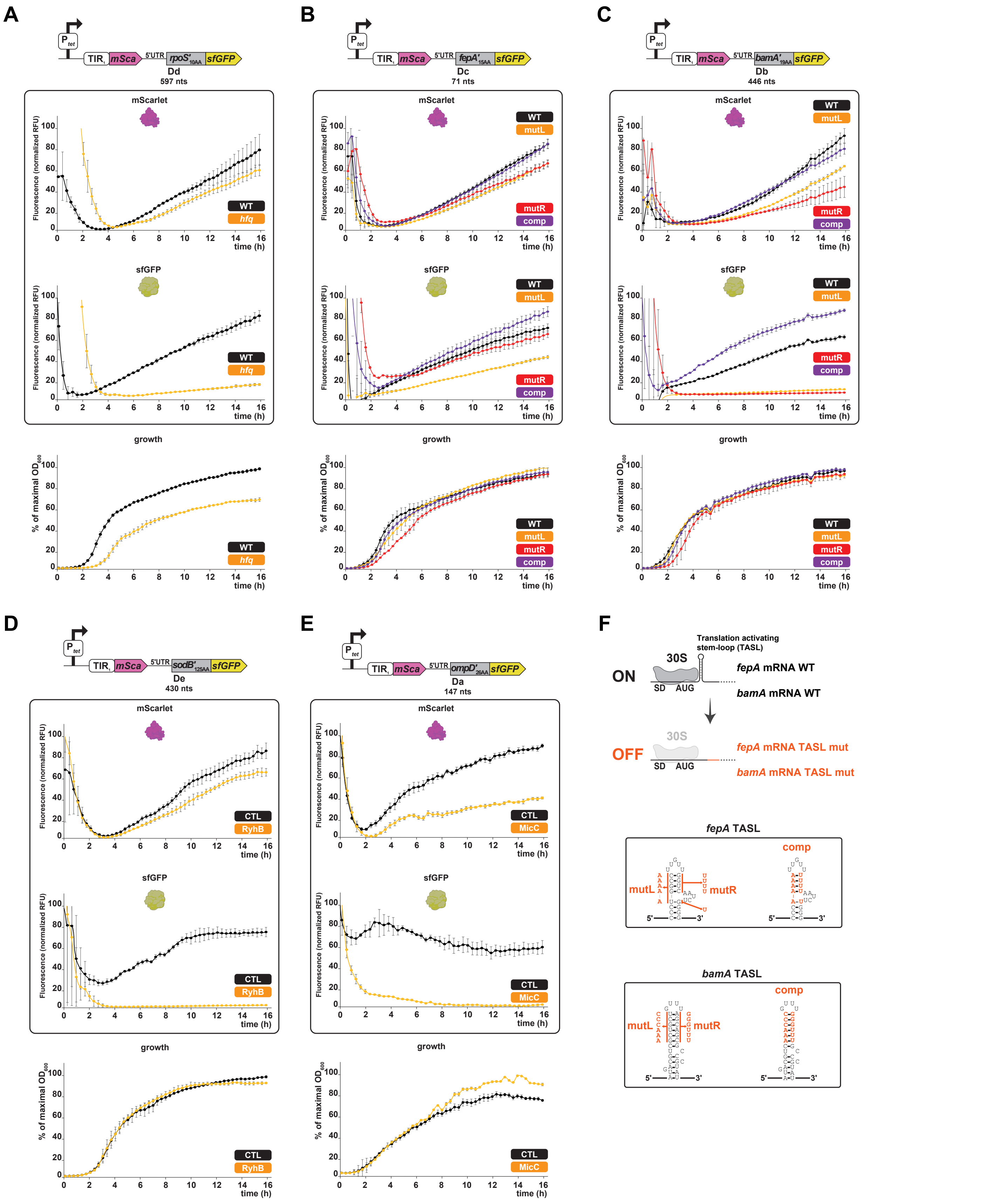


Figure S4. Fluorescence kinetics of the 5 tested genes in the tandem fluorescent reporters

A to E) Fluorescence kinetics were acquired for each strain as described in Fig. 5B using a microplate reader during 16h.

The examples of regulatory events affecting primarily translation tested in this work are the decrease of rpoS expression in an hfq deleted strain and the inhibition of the mRNA translation of bamA and fepA via the "mutL" mutations located in the left part of their TASL, as represented in Fig. S4F). Examples of regulatory events affecting mRNA stability tested in this work are the repression of sodB by overproduced sRNA RhyB and of ompD the overproduction of sRNA MicC respectively.

Strains carrying tandem fusions used are: hfq vs WT for rpoS: strains sML281 and sML280; TASL mutant (mutL, mutR and comp) vs WT for fepA: strains sML275, sML276, sML277 and sML274 and bamA: strains sML271, sML272, sML273 and sML270; sRNA expressing plasmid versus control plasmid for sodB: strains sML279 (pBR plac RyhB (11)) and sML278 (pBR plac (12)) and ompD: strains sML269 (pBR plac MicC (11))and sML268 (pBR plac (12)).

IPTG inducer of sRNA expression was added directly in the dilution medium at the beginning of growth. Fluorescence was measured in microplate reader for 16h. Fluorescence and growth kinetics are shown as mean relative fluorescence units (RFU on top and middle panels) and % of maximal OD600 (normalized homogeneously for both strains to the strain reaching the highest OD600, on the right). Values (Table S8) are means of 3 biological replicates and error bars are standard deviations.

Of note, some points are not shown in particular for early growth because we removed from the visualization of the data the measurements displaying negative fluorescence values and the measurements for which error bars were ≥10-fold higher than the maximal normalized fluorescence value. It should be noted that the hfq deletion strain shows a reduced growth rate as previously reported (13).

F) Schematic representation of the impact of introducing mutations that disrupt the translation activating stem loop in fepA or bamA on translation initiation (recruitment of the 30S subunit of the ribosome). The detailed mutations introduced in fepA and bamA are indicated on the bottom panels in red (mutL, mutR and comp).

### Figure S5

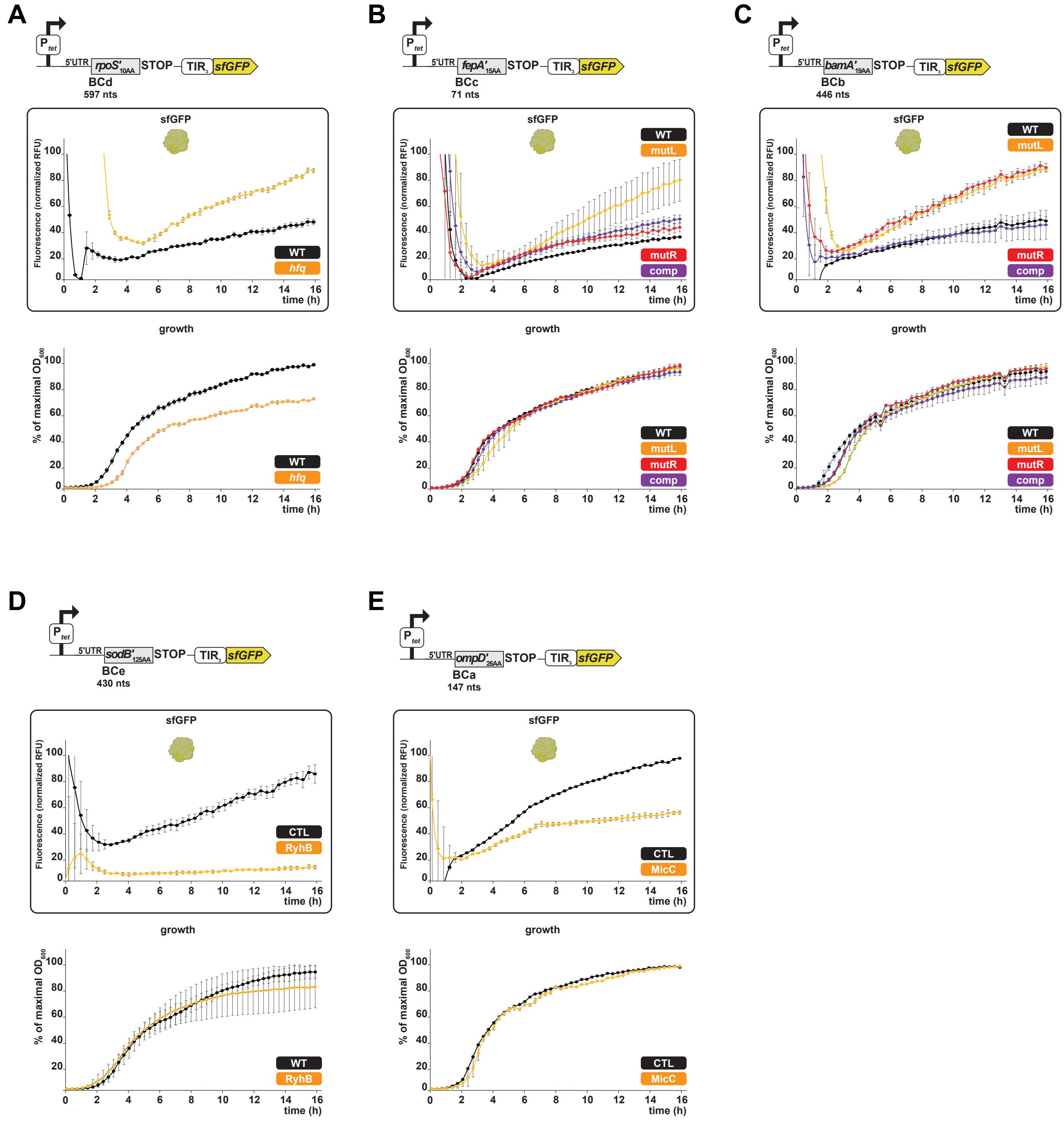


Figure S5. Fluorescence kinetic curve for 5 tested genes in the transcriptional reporter system

A to E) Fluorescence kinetics were acquired for each strain as described in Fig. 5C using a microplate reader during 16h.

Strains carrying mRNA read-out fusions used are: hfq vs WT for rpoS: strains sML295 and sML294; TASL mutant (mutL, mutR and comp) vs WT for fepA: strains sML289, sML290, sML291 and sML288 and bamA: strains sML285, sML286, sML287 and sML284; sRNA expressing plasmid versus control plasmid for sodB: strains sML293 (pBR plac RyhB (11)) and sML292 (pBR plac (12)) and ompD: strains sML283 (pBR plac MicC (11)) and sML282 (pBR plac (12)).

IPTG inducer of sRNA expression was added directly in the dilution medium at the beginning of growth. Fluorescence was measured in microplate reader for 16h. Fluorescence and growth kinetics are shown as mean relative fluorescence units (RFU on top and middle panels) and % of maximal OD600 (normalized homogeneously for both strains to the strain reaching the highest OD600, on the right). Values (Table S8) are means of 3 biological replicates and error bars are standard deviations.

Of note, some points are not shown in particular for early growth because we removed from the visualization of the data the measurements displaying negative fluorescence values and the measurements for which error bars were ≥10-fold higher than the maximal normalized fluorescence value.

### Figure S6

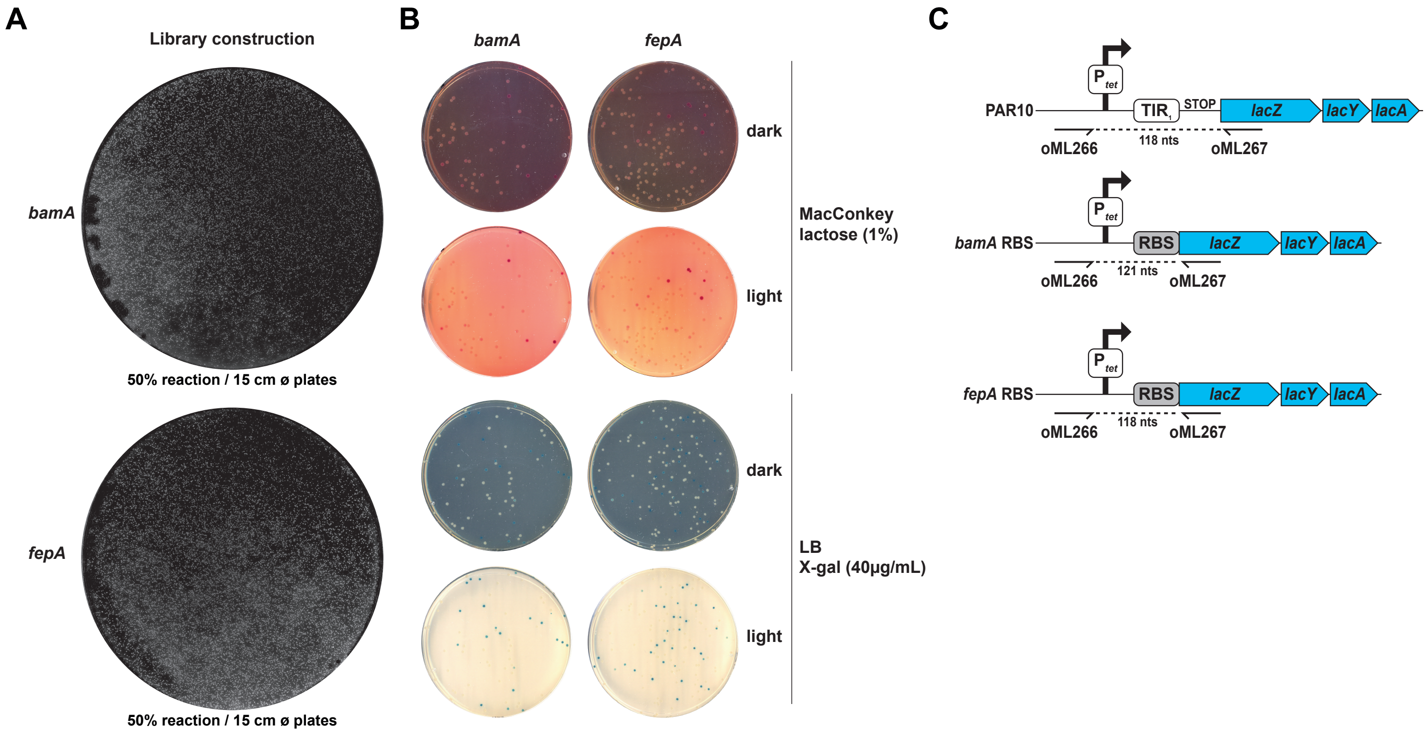


Figure S6. Coverage and phenotypic diversity in mutant libraries

A) Plated libraries (50% of the total reaction on a 15 cm diameter Petri dish) after editing are shown (white light with dark background scan).

B) Diluted individual libraries were plated on LB agar with X-gal or on MacConkey agar with lactose (white light with white or dark background scan).

C) Regions of complementarity of the PCR primers on the PAR10, bamA RBS and fepA RBS libraries construction are schematically represented. These regions were amplified from individual libraries and sent for deep-sequencing.
